# Tissue-resident FOLR2^+^ macrophages associate with tumor-infiltrating CD8^+^ T cells and with increased survival of breast cancer patients

**DOI:** 10.1101/2021.04.12.439412

**Authors:** Rodrigo Nalio Ramos, Yoann Missolo-Koussou, Yohan Gerber-Ferder, Christian Bromley, Mattia Bugatti, Nicolas Gonzalo Núñez, Jimena Boari Tosello, Wilfrid Richer, Jordan Denizeau, Christine Sedlik, Pamela Caudana, Fiorella Kotsias, Leticia Laura Niborski, Sophie Viel, Mylène Bohec, Sonia Lameiras, Sylvain Baulande, Laëtitia Lesage, André Nicolas, Didier Meseure, Anne Vincent-Salomon, Fabien Reyal, Charles-Antoine Dutertre, Florent Ginhoux, Lene Vimeux, Emmanuel Donnadieu, Bénédicte Buttard, Jérôme Galon, Santiago Zelenay, William Vermi, Pierre Guermonprez, Eliane Piaggio, Julie Helft

**Affiliations:** PSL University, Institut Curie Research Center, INSERM U932 & SiRIC, Center for cancers immunotherapy, Translational Immunotherapy Team, F-75005, Paris, France; The University of Manchester, CRUK Manchester Institute, Cancer Inflammation and Immunity Group, Manchester, UK; Department of Pathology, University of Brescia, Brescia 25123, Italy; Department of Pathology and Immunology, Washington University School of Medicine, St. Louis, MO 63110, USA; PSL University, Institut Curie Research Center, Institut Curie Genomics of Excellence (ICGex) Platform, F-75005, Paris, France; PSL University, Institut Curie Hospital, Department of Pathology, F-75005, Paris, France; PSL University, Institut Curie Hospital, Department of Surgery, F-75005, Paris, France; Paris-Saclay University, Institut Gustave Roussy, INSERM U1015, Villejuif, France; Singapore Immunology Network, Agency for Science, Technology and Research, Singapore 138648, Singapore; University of Paris, Institut Cochin, INSERM U1016, CNRS UMR 8104, F-75014 Paris, France; INSERM, Sorbonne Université, Université de Paris, Equipe Labellisée Ligue Contre le Cancer, Centre de Recherche des Cordeliers, Laboratory of Integrative Cancer Immunology, Paris, France; Université de Paris, Centre for Inflammation Research, CNRS ERL8252, INSERM1149 Paris, France

## Abstract

Macrophage infiltration is a hallmark of solid cancers and overall macrophage infiltration is correlated with lower patient survival and resistance to therapy. However, tumor-associated macrophages are phenotypically and functionally heterogeneous. Specific tumor-associated macrophage subsets might be endowed with antagonistic role on cancer progression and on the development of anti-tumor immunity. For instance, monocyte-derived TREM2^+^ tumor-associated macrophages have pro-tumorigenic and immunosuppressive functions. Here, we identify a discrete population of FOLR2^+^ tumor-associated macrophages positively correlating with patient survival in breast cancer. FOLR2^+^ macrophages are evolutionarily conserved across species and populate human and murine healthy mammary gland. Moreover, FOLR2^+^ macrophages co-localize with lymphoid aggregates containing CD8^+^ T cells in breast cancer and across ten other types of cancers. This study highlights antagonistic roles for tumor-associated macrophage subsets and paves the way for subset-specific therapeutic interventions in macrophages-based cancer therapies.

## INTRODUCTION

Macrophages are one of the most abundant immune cell population in human breast tumors microenvironment (TME)(Cassetta and Pollard, 2018). Macrophage infiltration in breast tumor correlates with poor prognosis and higher tumor grades (Zhao et al., 2017)(Ruffell and Coussens, 2015)(Ramos et al., 2020). Tumor-associated macrophages (TAMs) play pro-tumoral roles by providing growth factors to tumors, enhancing tumor cell motility and invasion, and by promoting angiogenesis and metastasis (Lewis and Pollard, 2006)(Engblom et al., 2016)(Caux et al., 2016). In addition, TAMs exert immunosuppressive functions thereby preventing tumor cell destruction by NK and T lymphocytes. Therefore, targeting TAM recruitment, survival and function has become a major therapeutic goals (Ries et al., 2014)(Mantovani et al., 2017)(Binnewies et al., 2018).

Even though the current paradigm presents TAMs as pro-tumorigenic cells in most instances, several studies have highlighted protective roles for TAMs in specific disease stages or organs (Ruffell and Coussens, 2015). Accordingly, targeting TAMs in preclinical cancer models can have a detrimental impact on tumor progression and metastasis (Bonapace et al., 2014)(Cassetta and Pollard, 2018)(Hanna et al., 2015). As described earlier (Mantovani et al., 2004), distinct populations of macrophages with opposite pro- and anti-tumoral functions might co-exist within the same tumor (Mantovani et al., 2002)(Ali et al., 2016). Therefore, establishing the extent of heterogeneity in the macrophage compartment is a pre-requisite for the rational design of macrophage-targeting therapies.

Macrophage heterogeneity might potentially arise from i) alternative activation states (Mantovani et al., 2017), ii) imprinting by tissue- or tumor-associated cues defining macrophage niches (Cassetta et al., 2019)(Guilliams and Scott, 2017); iii) distinct TAM cellular origins (adult monocyte versus embryonic progenitors)(Franklin et al., 2014)(Ginhoux et al., 2010)(Loyher et al., 2018)(Zhu et al., 2017) and iv) tumor-induced systemic modification of circulating monocytes (Gallina et al., 2006)(Veglia et al., 2018)(Cassetta et al., 2019)(Ramos et al., 2020).

In human breast cancer, macrophage infiltration has been assessed with markers like CD14 or CSF1R (Ruffell et al., 2012)(Cassetta et al., 2019) and CD68 (Leek et al., 1996)(Yuan et al., 2014). However, CD14 and CSF1R also mark undifferentiated monocytes while CD68 expression among phagocytes is not fully characterized. Other markers like CD163, TIE2, MRC1/CD206 or MARCO have been implemented to assess TAM phenotypic heterogeneity (Cassetta and Pollard, 2018). Pioneer single cell RNA sequencing (scRNAseq) studies have invalidated alternative activation as the main mechanism accounting for TAM heterogeneity (Azizi et al., 2018). In summary, the phenotypic and functional diversity of TAM infiltrating human breast cancer remain to be elucidated.

Here we implement scRNAseq of tumor associated CD14^+^HLA-DR^+^ cells isolated from metastatic LNs and primary breast tumors to assess the cellular heterogeneity within the CD14^+^ compartment. We identify two phenotypically distinct macrophage populations:

1. TREM2^+^ macrophages expressing Triggering Receptor Expressed by Myeloid cells-2 (*TREM2*), Osteopontin (*SPP1*) and Cell Adhesion Molecule 1 (*CADM1*) genes;
2. FOLR2^+^ macrophages expressing Folate Receptor 2 (FOLR2), Hyaluronan receptor (*LYVE-1*) and Mannose Receptor C-Type 1 (*MRC1/CD206*) genes.

We show that tumor-associated TREM2^+^ and FOLR2^+^ macrophages are evolutionarily conserved between human and mouse breast cancers. TREM2^+^ macrophages are poorly represented in healthy breast tissues but increase with tumor development. By contrast, we show that FOLR2^+^ macrophages are tissue-resident macrophages (TRMs) populating healthy mammary glands prior the onset of cancer development. Specific gene signatures defining FOLR2^+^ macrophages correlate with better relapse-free survival in breast cancer patients. Accordingly, FOLR2^+^ macrophages positively correlate with signatures of major cellular players of anti-tumor immunity, including CD8^+^ T cells, NK cells and dendritic cells (DCs). We further show that FOLR2^+^ macrophages locate near vessels and cluster with CD8^+^ T cell aggregates in tumor resection samples. This FOLR2^+^ macrophage/CD8^+^ T cell co-localization correlates with favorable clinical outcomes suggesting an anti-tumorigenic role for this newly characterized macrophage subset.

## RESULTS

### APOE expression defines tumor-associated macrophages in human breast cancer

Motivated by the idea of unambiguously identifying macrophages within breast tumors, we sought to define common features enabling the distinction between breast cancer macrophage populations and infiltrating CD14^+^CD1c^-^ monocytes, CD14^+^CD1c^+^ inflammatory DCs/DC3 or CD14^-^CD1c^+^ cDC2 (Villani et al., 2017)(Bourdely et al., 2020)(Dutertre et al., 2019).

In a first approach, we quantified mononuclear phagocytes of matched primary tumors, non-metastatic and metastatic lymph nodes (LNs) from a cohort of 13 treatment-naive luminal breast cancer patients (Table S1). We found that CD1c^-^CD14^+^ monocyte/macrophage population increased most significantly in metastatic LNs as compared to matched non-metastatic LNs (Fig. 1A). This positive correlation between CD14^+^ cell infiltration and tumor invasion, was confirmed in a second cohort of patients (Fig S1A). CD14^+^ cell infiltration correlated with the extent of tumor invasion in LNs (Fig. 1B).

**Figure 1.**
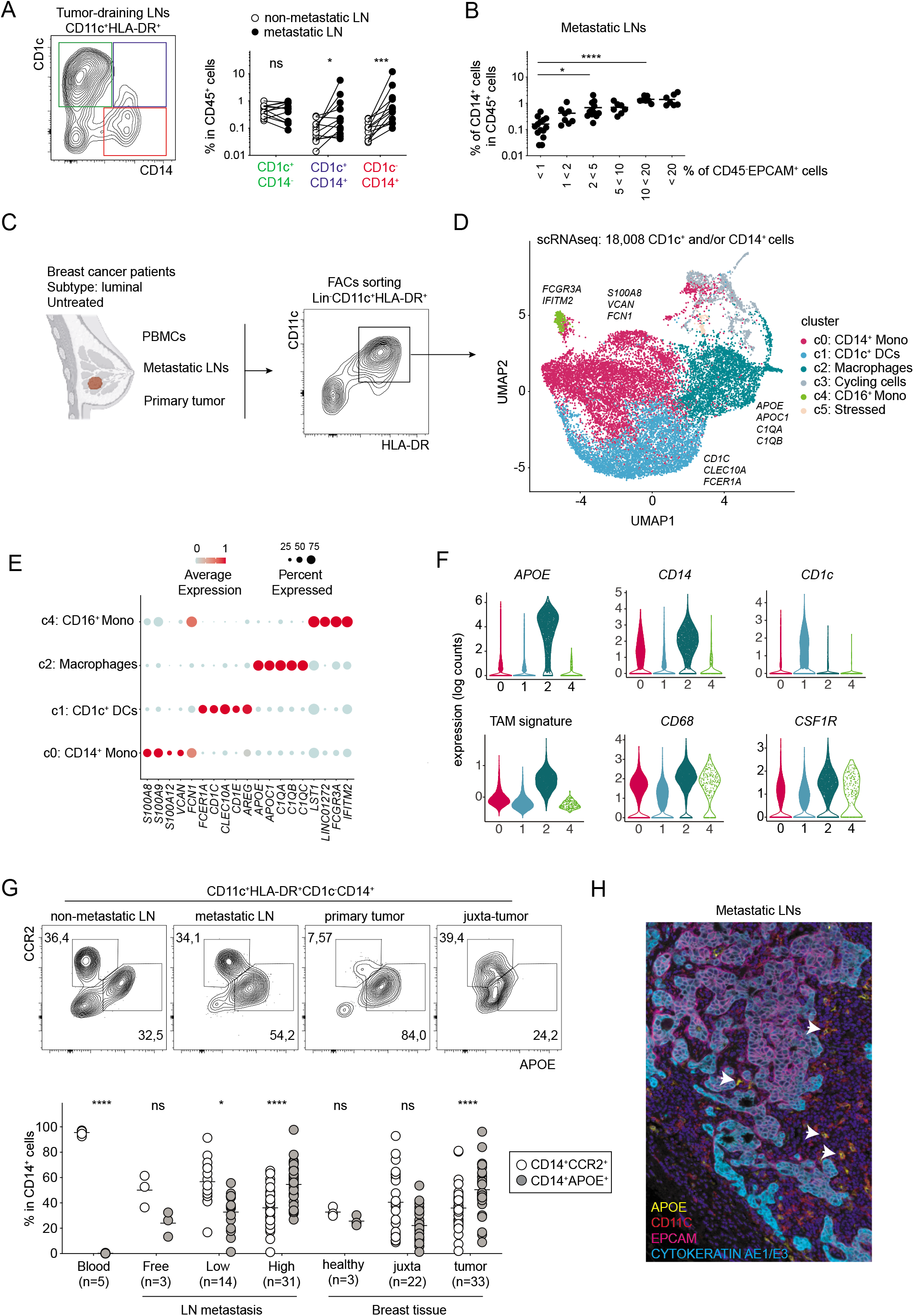
APOE expression defines tumor-associated macrophages in human breast cancer. **A.** Flow cytometry quantification of CD1c^+^CD14^-^, CD1c^+^CD14^+^ and CD1c^-^CD14^+^ cells among CD11c^+^HLA-DR^+^ myeloid cells in matched non-metastatic and metastatic lymph nodes. **B.** Flow cytometry quantification of CD14^+^ cells in stratified metastatic lymph nodes according to CD45^-^EPCAM^+^ tumor cells frequency. **C.** Experimental design of untreated luminal breast cancer patient samples (PBMCs, metastatic lymph nodes and primary tumors) and gating strategy for FACS-sorting of myeloid cells for scRNAseq droplet-based assay (10x Chromium). **D.** Dimensionality reduction of scRNAseq data merged from blood, metastatic lymph nodes and primary tumors was performed using a Louvain graph-based clustering identifying six clusters. Each dot represents an individual cell (n=18008). Representative genes from each cluster are depicted. **E.** Bubble-map showing the top five most significant expressed genes across the six defined clusters. Circles sizes and represent percentage of cells within a cluster expressing a gene and color represents the average expression of each gene. **F.** Violin plots illustrating expression distributions of selected genes of interest in clusters 0 (CD14^+^ monocytes), cluster 1 (CD1C^+^ DCs), cluster 2 (macrophages) and cluster 4 (CD16^+^ monocytes). **G.** Flow cytometry quantification of CD14^+^CCR2^+^ monocytes versus CD14^+^APOE^+^ macrophages in healthy or metastatic lymph nodes and breasts. (*P≤ 0.05; **P≤ 0.01; ****P< 0.0001). **H.** Immunofluorescence images of APOE^+^ macrophages and EPCAM^+^Cytokeratin^+^ tumor cells in a representative metastatic lymph nodes.

We next sought to characterize the heterogeneity within the entire tumor-infiltrating CD14^+^ cells in an unbiased manner. To this end, we analyzed metastatic LNs, primary tumors and blood of untreated luminal breast cancer patients (Fig. 1C, Table S1). We isolated all mononuclear phagocytes by FACs-sorting the CD11c^+^HLA-DR^+^ cell fraction and performing scRNAseq using a droplet-based approach. We used the SEURAT pipeline to process the data (Fig. 1D). We excluded from the analysis: contaminating lymphocytes (B/T/NK cells), XCR1^+^ DCs and CCR7^+^LAMP3^+^ DCs (Zhang et al., 2019). We then merged the ~ 18000 remaining myeloid cells from all the patients (Fig. 1D, S1B). Louvain Graph-based clustering identified 4 clusters of mononuclear phagocytes and populations of cycling (*mKI67, TOP2A, CDC20, e.g.*) and “stressed” cells (*HSPA1A, HSPB1, e.g.*) (Fig S1C, Table S2). Cluster 0 was characterized by the selective and high expression of markers (*S100A8, S100A9, S100A12, VCAN*) defining CD14^+^CD16^-^ monocytes (Villani et al., 2017)(Fig. 1D, E, Table 2). Cluster 1 was characterized by genes defining CD1c^+^ DCs (including DC2 and DC3 subsets) while cluster 4 was identified as CD14^-^CD16^+^ monocytes (Villani et al., 2017)(Dutertre et al., 2019)(Bourdely et al., 2020). Monocyte-clusters (c0 and c4) were both found in blood, tumor and metastatic LNs (Fig S1C). The remaining cluster 2 was identified as TAMs because it selectively expressed high levels of a TAM signature (Fig. 1F)(Azizi et al., 2018). Cluster 2 expressed high levels of *APOE, APOC1, C1QA, C1QC* enabling the distinction from monocytes (Fig. 1E, S1D). We show that homogenous expression of APOE selectively discriminate TAMs from both CD14^+^ monocytes and CD1c^+^DCs (Fig. 1F). No other commonly used markers (CSF1R, CD68, CD14) achieved this discrimination. CD68 is expressed in CD14^+^ monocytes, CD16^+^ monocytes and CD1c DCs; CD14 is expressed by monocytes and a subset of CD1c^+^ DCs; CSF1R is promiscuous (Fig. 1F).

We next wanted to validate protein expression distinguishing macrophages/TAMs from monocytes within CD1c^-^CD14^+^ cells. The intracellular proteins S100A8 and S100A9 - found to be highly and specifically expressed in CD14^+^ monocyte cluster 0 - followed the protein-expression pattern of cell-surface CCR2, which was not detected in our scRNAseq dataset (Fig S1E). The best monocyte/macrophage discrimination was obtained by staining with APOE and CCR2 (Fig. 1G). We next analyzed the contribution of CCR2^+^ monocytes versus APOE^+^ macrophages to the CD14^+^ cell compartment within luminal breast tumor lesions (primary tumor or highly invaded metastatic LNs) versus tumor-free tissues (blood, healthy breast tissue, tumor-free and lowly invaded metastatic LNs or juxta-tumor). We found that the frequency of APOE^+^ macrophages increased with tumor burden while the frequency of CCR2^+^ monocytes decreased (Fig. 1G). Finally, we found that APOE^+^ cells located near and inside the tumor lesions in metastatic LNs and primary tumors (Fig. 1H, S1F). Altogether, our results establish APOE as a specific marker to identify macrophages in primary and metastatic luminal breast cancer.

### Single cell RNA sequencing reveals two main subsets of APOE^+^ macrophages

We next investigated the heterogeneity of APOE^+^ TAMs (cluster 2). Louvain graph-based clustering identified 3 clusters of TAMs (Fig. 2A, S2A-B). Hierarchical clustering showed that clusters 0 and 1 were closer to each other as compared to cluster 2 (Fig. 2B). To further explore the transcriptional heterogeneity found within the APOE^+^ macrophages, we implemented single-cell regulatory network inference and clustering (SCENIC) to study the gene regulatory network (GRN) of each macrophage cluster. GRN analysis and their activity-based hierarchical clustering revealed that cluster 0 and 1 shared around half of their transcriptional regulon including CEBPB and BHLHE41 while cluster 2 presented mostly unique transcriptional regulons like NR1H3 and MAF (Fig. 2C; Table S3). Analysis of differentially expressed genes revealed that *FOLR2, SEPP1, SLC40A1, MRC1, LYVE1*, discriminated cluster 2 from both cluster 0 and cluster 1 (Fig. 2D; Table S4). Conversely, *TREM2, SPP1, ISG15* discriminated both cluster 0 and cluster 1 from cluster 2 (Fig. 2D; Table S4). Altogether, we show that *FOLR2* is a defining marker for cluster 2 while *TREM2* defines cluster 0 and 1 (Fig. 2E). We conclude that APOE^+^ TAMs comprise two distinct populations: the TREM2^+^ macrophages and the FOLR2^+^ macrophages.

**Figure 2.**
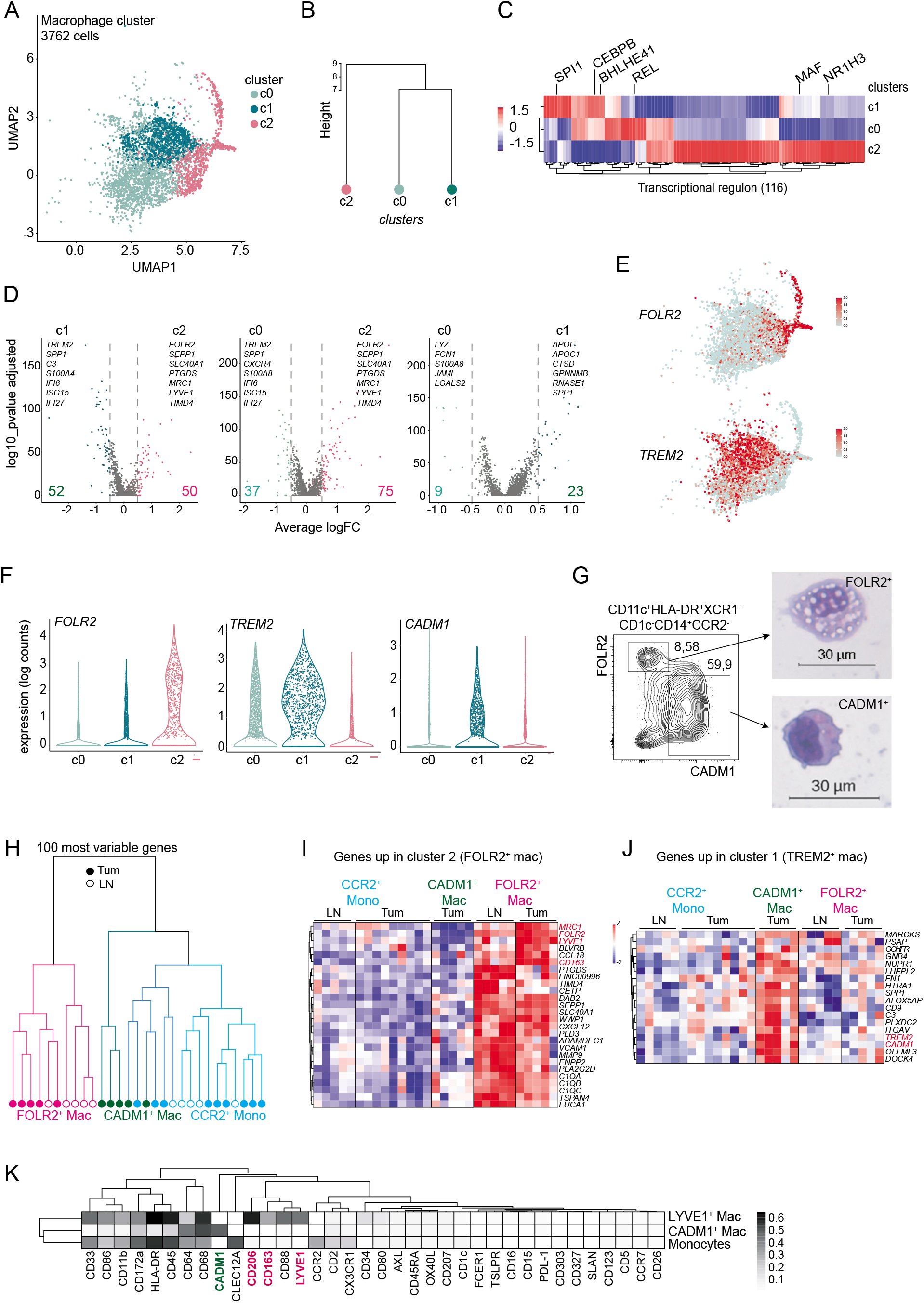
Single cell RNA sequencing reveals two main subsets of APOE^+^ macrophages. **A.** UMAP plot visualization of APOE^+^ macrophages (cluster 2 from Fig 1D). Each dot represents an individual cell (n=3762) **B**. Hierarchical clustering of clusters 0, 1 and 2 based on average gene expression (1200 genes). **C.** Heatmap and hierarchical clustering of differently predicted transcriptional regulons in clusters 0, 1 and 2 defined using the SCENIC pipeline. **D.** Volcano plot showing D.E.G. between each cluster. Selected genes among the Top25 were depicted. **E.** UMAP plot showing mutually exclusive expression of *TREM2* and *FOLR2* in *APOE^+^* macrophages. **F.** Violin plots expression distributions of *FOLR2*, *TREM2* and *CADM1* across *APOE*^+^ macrophage clusters **G.** Representative flow cytometry contour-plot and cytospin images from FACS-sorted FOLR2^+^ and CADM1^+^ macrophages isolated from metastatic lymph nodes and primary tumors. **H**. Hierarchical clustering using the 100 most variable genes from bulk RNA sequencing of FOLR2^+^ TAMs, CADM1^+^ TAMs and CCR2^+^ monocytes isolated by FACS-sorting from metastatic lymph nodes and primary tumor of untreated luminal breast cancer patients. **I-J.** Heatmap of the D.E.G. between *FOLR2*^+^ (c2) and *TREM2*^high^ (c1) macrophages selected from scRNAseq dataset and applied to bulk RNAseq of FOLR2^+^ macrophages, CADM1^+^ macrophages and CCR2^+^ monocytes isolated by FACS-sorting from metastatic lymph nodes and primary tumor of untreated luminal breast cancer patients. **K.** Protein expression analysis (CyTOF) of monocyte/macrophage and dendritic cell markers in CD14^+^CCR2^+^ monocytes, CD14^+^CCR2^-^CD68^+^LYVE1^+^ macrophages and CD14^+^CCR2^-^CD68^+^CADM1^+^ macrophages from metastatic lymph nodes.

We next sought to validate this finding by prospective isolation of these populations for bulk transcriptome analysis. To this end, we searched for surface proteins differentially expressed between the two populations thus enabling their isolation by flow cytometry. We failed to detect TREM2 at the cell surface of TAMs after tissue dissociation. Alternatively, we found that cluster 1 (TREM2^high^ TAMs) specifically expressed and stained positive for CADM1 (Fig. 2F, G).

Flow cytometry analysis of FOLR2 and CADM1 expressions revealed mutually exclusive expression patterns (Fig. 2G). We isolated FOLR2^+^CADM1^-^ and FOLR2^low^CADM1^+^ TAMs from both primary tumors and metastatic LNs by FACs sorting (Fig S2D). FOLR2^+^CADM1^-^ TAMs presented a typical macrophage shape and were filled with vacuoles. In contrast, FOLR2^low^CADM1^+^ TAMs were smaller in size, with a morphology closer to monocytes (Fig. 2G).

We next performed bulk RNAseq on FOLR2^+^CADM1^-^ TAMs, FOLR2^low^CADM1^+^ TAMs and CD14^+^CCR2^+^ monocytes (Fig. 2H, Table S1). Hierarchical clustering showed that FOLR2^+^CADM1^-^ macrophages from both primary tumors and invaded LNs cluster together away from FOLR2^low^CADM1^+^ macrophages or CD14^+^CCR2^+^ monocytes (Fig. 2H). We confirmed our scRNAseq results showing that FOLR2^+^ macrophages expressed higher levels of *FOLR2, SEPP1, SLC40A1* and *LYVE1* (Fig. 2I) as compared to FOLR2^low^CADM1^+^ TAMs and CD14^+^CCR2^+^ monocytes. Overall, FOLR2^low^CADM1^+^ macrophages from primary tumors clustered together with CD14^+^CCR2^+^ monocytes (Fig. 2H). However, FOLR2^low^CADM1^+^ macrophages specifically expressed *TREM2* and genes found to be overexpressed in cluster 1 (*C3, FN1, SPP1*) of the scRNAseq analysis (Fig. 2J).

Some of these phenotypic differences were confirmed by CyTOF profiling of CD14^+^CCR2^-^ macrophages from invaded LNs. Co-expression of LYVE1, MRC1/CD206 and CD163 was found in a macrophage population distinct from CADM1^+^ expressing macrophages (Fig. 2K).

Altogether, our results show that breast TAMs comprise two populations separable by their mutually exclusive expression of TREM2/CADM1 and FOLR2.

### FOLR2^+^ macrophages are tissue-resident macrophages

A recent study identifies the infiltration of TREM2^+^ macrophages as an event associated to cancer development (Molgora et al., 2020). Our data establish their transcriptional proximity to CD14^+^CCR2^+^ monocytes in breast tumors (Fig. 2H). These results suggest that TREM2^+^CADM1^+^ TAMs arise from infiltration of circulating monocytes during tumor progression.

The origin of FOLR2^+^ macrophage is not known. We wondered whether FOLR2^+^ macrophages correspond to mammary TRMs (*i. e*. present in healthy breast) or tumor-recruited monocyte-derived macrophages like the TREM2^+^ TAMs. To address this question, we quantified FOLR2^+^ macrophages by flow cytometry in healthy tissues (healthy breast, mammary tissues adjacent to tumor lesion-juxta-tumor-, tumor-free or lowly-invaded metastatic LNs) versus luminal breast tumor lesion (primary tumors and highly invaded metastatic LNs). We found that among APOE^+^ macrophages, FOLR2^+^ macrophages were enriched in healthy and juxta-tumor tissues (Fig. 3A). By contrast, the fraction of FOLR2^-^ TAMs (comprising TREM2^+^ TAMs) dramatically increased in both primary and metastatic tumor lesions as compared to healthy tissues (Fig. 3A).

**Figure 3.**
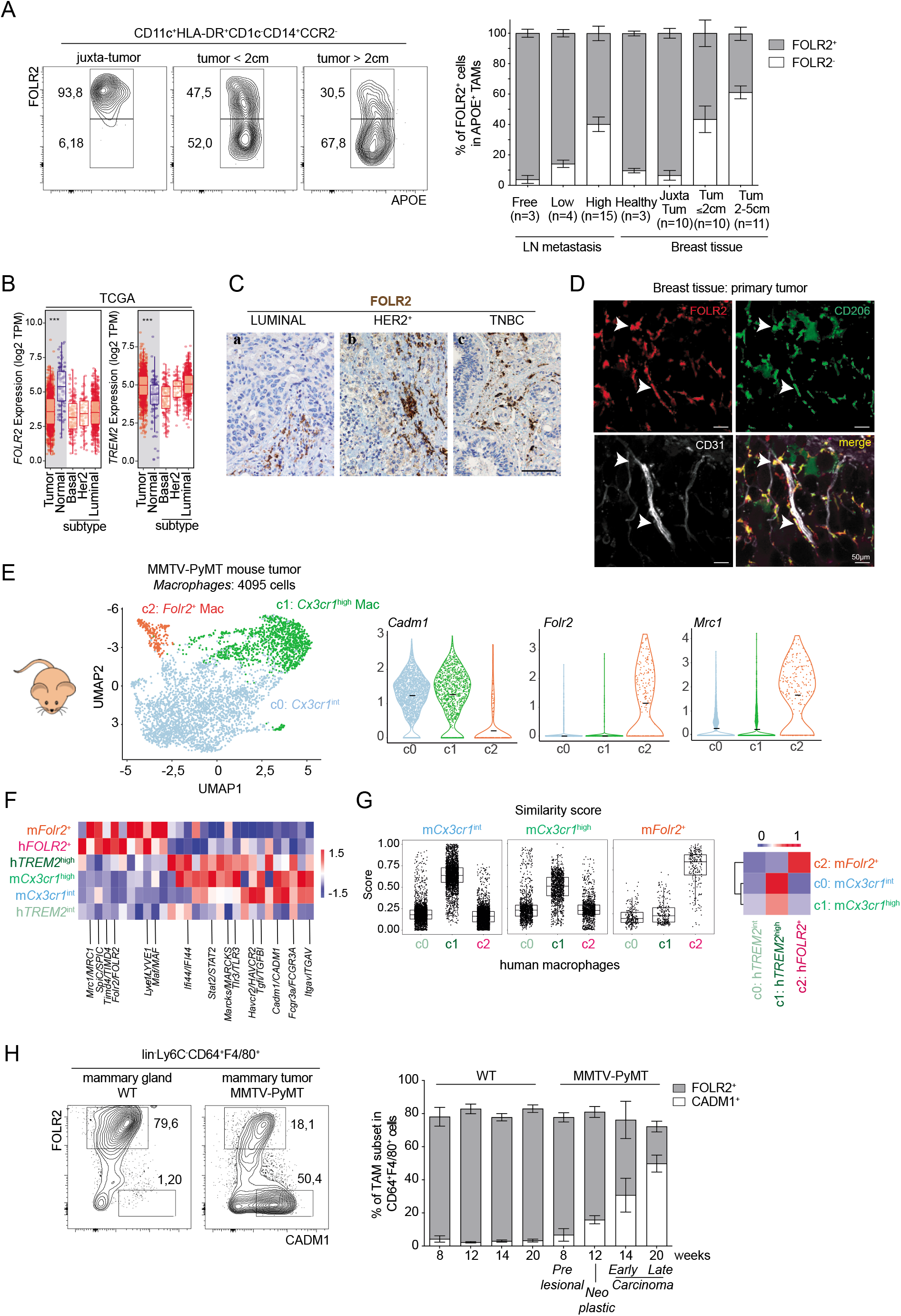
FOLR2^+^ macrophages are tissue-resident macrophages. **A.** Representative contour-plots and quantification of FOLR2^+^APOE^+^ and FOLR2^-^APOE^+^ macrophages across distinct breast cancer patient tissues using flow cytometry. **B.** *FOLR2* and *TREM2* mRNA expression in tumor microarrays from breast cancer patients of the TCGA database (Tumor: n=1093; Normal breast: n=112; Basal subtype, n=190; Her2 subtype, n=82; and Luminal subtype, n=781). **C.** FOLR2 expression by immunohistochemistry. Sections are from three cases of primary breast carcinomas showing a luminal (**a**), a HER2^+^ (**b**) and a triple negative (**c**) subtype. Sections are counterstained with Mayer’s hematoxilin. Magnification=200X, scale bar 100 micron. **D.** Representative confocal immunofluorescence images performed in primary breast cancer tissues. White arrows indicate FOLR2^+^CD206^+^ macrophages co-localizing with CD31^+^ vessels. **E.** UMAP plot visualization of *Fcgr1* macrophages (n= 4095) isolated from mammary tumors of 23-week-old MMTV-PyMT mice (n=2). Violin plots illustrating expression distributions of *Cadm1, Folr2 and Mrc1* across *Fcgr1^+^* macrophages. **F.** Heatmap showing gene orthologs similarly expressed across mouse and human macrophage subsets. **G.** Similarity score defined using the Seurat v3 reference label transfer integration (Stuart et al., 2019). Graph plots showing prediction scores of each mouse macrophage cluster (defined in Fig. 3D) applied to each human macrophage cluster (defined in Fig. 2A). Each dot represents a cell. Heatmap of the mean of the prediction score. **H.** Quantification of Folr2^+^ and Cadm1^+^ macrophages by flow cytometry during tumor development in MMTV-PyMT mouse model. Upper part shows representative contour-plots of macrophage subsets in mammary gland from tumor-free and tumor-bearing mice (20-week-old PyMT mouse). Lower part shows quantification of macrophage subsets in WT or PyMT mice during tumor development.

The enrichment of FOLR2^+^ macrophages in adjacent normal tissue versus tumor lesions was also confirmed at the transcriptional level by analyzing breast cancer samples from the Cancer Genome Atlas (TCGA) database (Fig. 3B). *FOLR2* transcripts were enriched in normal adjacent tissues as compared to breast cancer tumor lesions of different subtypes (Her2^+^, TNBC, Luminal). *FOLR2* transcripts were also enriched in non-disease healthy tissues as compared to tumor in breast cancer (Fig S3A). In contrast *TREM2* transcripts were enriched in breast tumor lesions as compared to tumor-adjacent normal and non-disease healthy tissues (Fig. 3B, S3A). Importantly, immunohistochemistry (IHC) analysis revealed that FOLR2^+^ macrophages were present in peri-tumoral areas in all subtypes of breast cancers (Fig. 3C). Bulk RNAseq (Fig. 2J) and CyTOF (Fig. 2K) analysis of FOLR2^+^ macrophages show that FOLR2^+^ macrophages specifically express the hyaluronan receptor LYVE1 and the mannose receptor (MRC1/CD206), both markers of perivascular macrophages (Lin et al., 2006)(Lim et al., 2018)(Chakarov et al., 2019). Therefore, we wondered whether FOLR2^+^ macrophages would locate near vessels. Confocal imaging on tumor surgical resection specimens showed that FOLR2^+^CD206^+^ macrophages located near CD31^+^ vessels in both tumor and adjacent tissue (Fig. 3D, S3B). Altogether these results show that FOLR2^+^ macrophages are perivascular TRMs associated with healthy mammary glands and LNs.

We next analyzed mammary gland macrophages in the mouse model enabling us to carry out a longitudinal analysis of immune populations in steady state and during tumor progression. We first analyzed a published scRNAseq dataset performed on hematopoietic cells from healthy mammary glands (Han et al., 2018). We identified a subset of TRMs co-expressing *Folr2, Mrc1* and *Lyve1*, like human FOLR2^+^ macrophages (Fig S3C). These cells align to previously described MRC1^+^LYVE1^+^TRMs (Franklin et al., 2014)(Jäppinen et al., 2019)(Wang et al., 2020). We next performed scRNAseq on CD45^+^CD3^-^CD19^-^B220^-^NKP46^-^ cells isolated from MMTV-PyMT autochthonous luminal-like mammary tumor model (Franklin et al., 2014)(Davie et al., 2007)(Fig 3E, Fig S3D). We excluded from the analysis contaminating lymphocytes (not shown), *Ly6c2^+^* monocytes (c3), *Ly6c2^-^Nr4a1*^high^ monocytes (c6), cycling cells (c4) and cells with high content of ribosomal genes (c1)(Fig. S3D). Among the remaining *Fcgr1^+^* cells, we identified 3 clusters of TAMs in 23 weeks-old PyMT mouse-tumors (Fig. 3E). Among these, two expressed *Cadm1* (*Cx3cr1*^int^ cluster 0 and *Cx3cr1*^high^ cluster 1, Fig. 3E). In addition, we identified a discrete population of *Folr2^+^Mrc1^+^* macrophages (cluster 2, Fig. 3E). Mouse and human FOLR2^+^ TAMs share the expression of *FOLR2, MRC1, SLC40A1 and MAF* (Fig. 3F; Table 5). On the other hand, *Cadm1^+^Cx3cr1^+^* mouse macrophages (clusters 0 and 1) resemble human CADM1^+^TREM2^high^ TAMs (cluster 1, Fig. 2A) and share the expression of *CADM1, HAVCR2, IFI44* (Fig. 3F; Table 5). Of note, the *Trem2* expression pattern is more conspicuous in murine as compared to human macrophages (Fig. S3E). To probe the similarity between human and mouse FOLR2^+^ macrophages, we performed a similarity analysis across whole transcriptomes at the level of each cell (Fig. 3G). This unbiased analysis confirmed the marker-based alignment of murine *Folr2^+^* macrophage to human *FOLR2^+^* macrophages. Conversely, the *Cadm1^+^Cx3cr1^+^* murine macrophages (clusters 0 and 1) presented high similarity with CADM1^+^TREM2^+^ human macrophages. Thus, we concluded that FOLR2^+^ macrophages are evolutionarily conserved between murine and human luminal mammary tumors.

We next analyzed longitudinally the dynamics of FOLR2^+^ (and CADM1^+^) macrophages during the development of murine breast cancer. To this end, we analyzed mammary gland macrophages in healthy littermate (WT), pre-lesion PyMT mice (8 weeks-old), neoplastic lesions (12 weeks-old mice), early carcinoma (14 weeks-old mice) and advanced carcinoma (20 weeks-old mice). We found that FOLR2^+^ macrophages constitute around 90% of total macrophages (CD45^+^Lin^-^Ly6C^-^F4/80^+^CD64^+^), in healthy mammary gland (WT)(Fig S3F). The frequency of FOLR2^+^ macrophages progressively decreased upon carcinoma progression to reach a minimum of 10-20% in advanced-carcinoma lesion of 20 weeks-old PyMT mice (Fig. 3H). In contrast, carcinoma development was accompanied by the *de novo* expansion of CADM1^+^ macrophages representing up to 80% of total macrophage in 20 weeks-old PyMT mice (Fig. 3H). Altogether we concluded that FOLR2^+^ macrophages represent an evolutionarily conserved TRM subset persisting in advanced carcinoma.

### FOLR2^+^ macrophages correlate with increased survival in breast cancer patients

TAMs are generally thought to promote tumor growth and inhibit anti-tumor immunity. This is particularly well established in mouse models where macrophages display a plethora of pro-tumoral function (Lin et al., 2006)(Qian et al., 2011)(Franklin et al., 2014)(Linde et al., 2018). In human, TAMs generally correlate with poor prognosis and higher tumor grade. However, clinical studies have probed the association of TAMs to patient survival by using markers often shared by monocytes and macrophages (*CSF1R, MRC1/CD206, CD163* e.g) or even shared with other immune cells types like pDCs (*CD68* e.g.) (Colonna et al., 2004). Therefore, we wondered whether FOLR2^+^ macrophages are similarly associated with worse survival in breast cancer patients.

To this end, we defined gene signatures enabling us to infer the abundance of total macrophages or FOLR2^+^ macrophage subset within bulk tumor RNA sequencing. Three genes (*C1QA, C1QB, C1QC*) define a core macrophage signature shared by the 3 macrophage clusters identified in this study (Fig. 2A, Fig. 4A) and suffice to distinguish macrophages from other leukocytes lineages (Fig. 4A, B; Table 6). Three genes (*FOLR2, SEPP1, SLC40A1*) uniquely distinguish FOLR2^+^ macrophages from other TAMs and other leukocytes lineages (Fig. 4A, B)(Azizi et al., 2018)(Table S6). LYVE1 was not considered for the signature analysis because of its endothelial expression.

**Figure 4.**
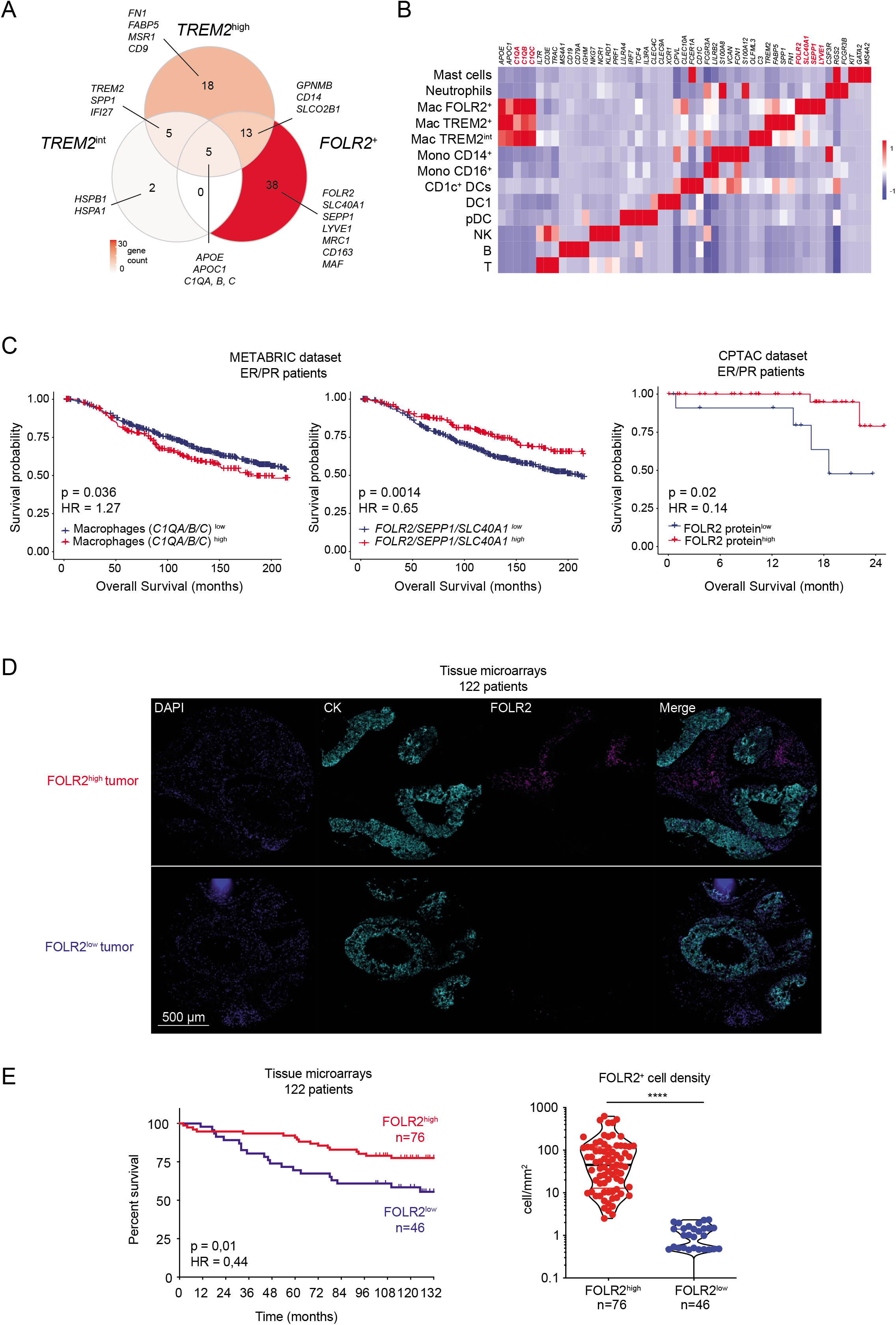
FOLR2^+^ macrophages correlate with increased survival in breast cancer patients. **A.** Venn diagram showing specific and common differentially expressed genes (DEG) of each APOE^+^ macrophages clusters (defined in Fig 2A). **B.** Heatmap showing mean expression of specific genes discriminating tumor-infiltrating immune cell populations in a published scRNAseq dataset performed on CD45^+^ cells isolated from breast cancer patients (Azizi et al., 2018). **C.** Kaplan-Meier survival curves generated for a macrophage gene signature (*C1QA/C1QB/C1QC*) and a FOLR2^+^ TAM gene signature (*FOLR2/SEPP1/SLC40A1*) in the METABRIC luminal breast cancer cohort (n=1309). Kaplan-Meier survival curve generated for FOLR2 protein expression in the CPTAC luminal breast cancer cohort (n=49). Patients were divided in high- and low-expressing groups based on 75% quantile of signature expression. **D.** Representative images of multiplex immunofluorescence from tissue-microarray showing FOLR2^high^ (top) and FOLR2^low^ (bottom) profiles. **E.** Kaplan-Meier survival curves generated for FOLR2^+^ macrophage density calculated by multispectral analysis of tumors (n=122). Graph shows the quantification of FOLR2^+^ macrophage density in tumors. Patients were divided in high- and low-cell-density groups based on best p value cut-off (**P≤0.01).

We analyzed the representation of these signatures within bulk transcriptomes of luminal breast cancer (1339 patients, METABRIC cohort)(METABRIC Group et al., 2012). Survival probability was calculated within highest signature expressers (top 25%) versus the rest of the patients (Fig. 4C). In accordance with previous reports (Ruffell and Coussens, 2015), we found that the highest level of macrophage infiltration correlated with worse overall survival (p=0,036). In stark contrast, highest level of FOLR2 signature correlated with increased overall survival (p=0,0013)(Fig. 4C). We also analyzed the association of FOLR2 protein and patient prognosis within 49 ER^+^/HER2^-^ patients from the CPTAC dataset. We found that FOLR2 protein abundance positively correlated with better survival (Fig 4C). Since FOLR2^+^ macrophages are a tissue-resident population in healthy mammary gland, *FOLR2* mRNA abundance could be associated to smaller and less aggressive tumors. To test whether this could be a confounding factor in the univariate survival analysis, we analyzed the level of expression of *FOLR2* mRNA for breast tumors of different stages and grade. We found no significant differences in *FOLR2* expression between grades and a slight increase in late stage tumors (Fig S4A). Moreover, multivariate analysis of the prognostic value of the FOLR2 gene-signature adjusted for age, histological grade, tumor size, histology and number of disease-positive LNs, showed that the FOLR2 gene-signature was an independent prognostic factor correlating with better survival of luminal breast cancer patients (Fig S4B).

We next wanted to directly assess if the density of FOLR2^+^ macrophages was associated with favorable clinical outcomes. To this end, we used multispectral imaging to analyze a tissue microarray comprising tumors from a retrospective cohort of 122 breast cancer patients. We stained the tumors for FOLR2, cytokeratin and DAPI and calculated the cellular density of FOLR2^+^ macrophages (Fig. 4D). Using the best performing threshold as a cut-off, we found that FOLR2^+^ macrophage density positively correlates with patient survival (Fig. 4E). Altogether these results show that FOLR2^+^ macrophage-abundance is associated with better prognosis for breast cancer patients.

### FOLR2^+^ macrophages are spatially segregated from TREM2^+^ macrophages across cancers

TREM2^+^ TAMs have been shown to infiltrate tumor nest across cancers (Molgora et al., 2020). We have shown that FOLR2^+^ macrophages are mammary-gland TRMs. Moreover, others have recently shown that macrophages expressing FOLR2 are found in healthy human tissues (Samaniego et al., 2014) (Sharma et al., 2020)(Thomas et al., 2021). Therefore, we wondered whether we could detect FOLR2^+^ macrophages across cancer types. To address this question, we analyzed FOLR2 expression in 80 histological sections of primary and metastatic tumors across distinct cancers (oral cavity, liver, bladder, brain, kidney, skin, colon, lung, ovary, stomach, breast)(Table S7). We found FOLR2^+^ macrophages across all these cancers (Fig. 5A, S5). Co-staining for FOLR2 and TREM2 confirmed mutually exclusive expression of the two markers on distinct cells (Fig. 5B). Staining of FOLR2 and TREM2 on serial sections of various cancer types showed that FOLR2^+^ and TREM2^+^ macrophages are spatially segregated. As described by Molgora *et al*, TREM2^+^ macrophages infiltrated the tumor nest (Fig. 5A). By contrast, FOLR2^+^ macrophages were consistently found within peri-tumoral stromal areas (Fig. 5A). This spatial segregation was found in various cancer types including melanoma (Fig 5Aa), lung carcinoma (Fig 5Ab, c), hepatocellular carcinoma (Fig 5Ad), oral cavity squamous cell carcinoma (Fig 5Ae) and pancreatic carcinoma (Fig 5Af). Altogether, these results show that FOLR2^+^ and TREM2^+^ macrophages are associated to specific location within the TME in various cancer types.

**Figure 5.**
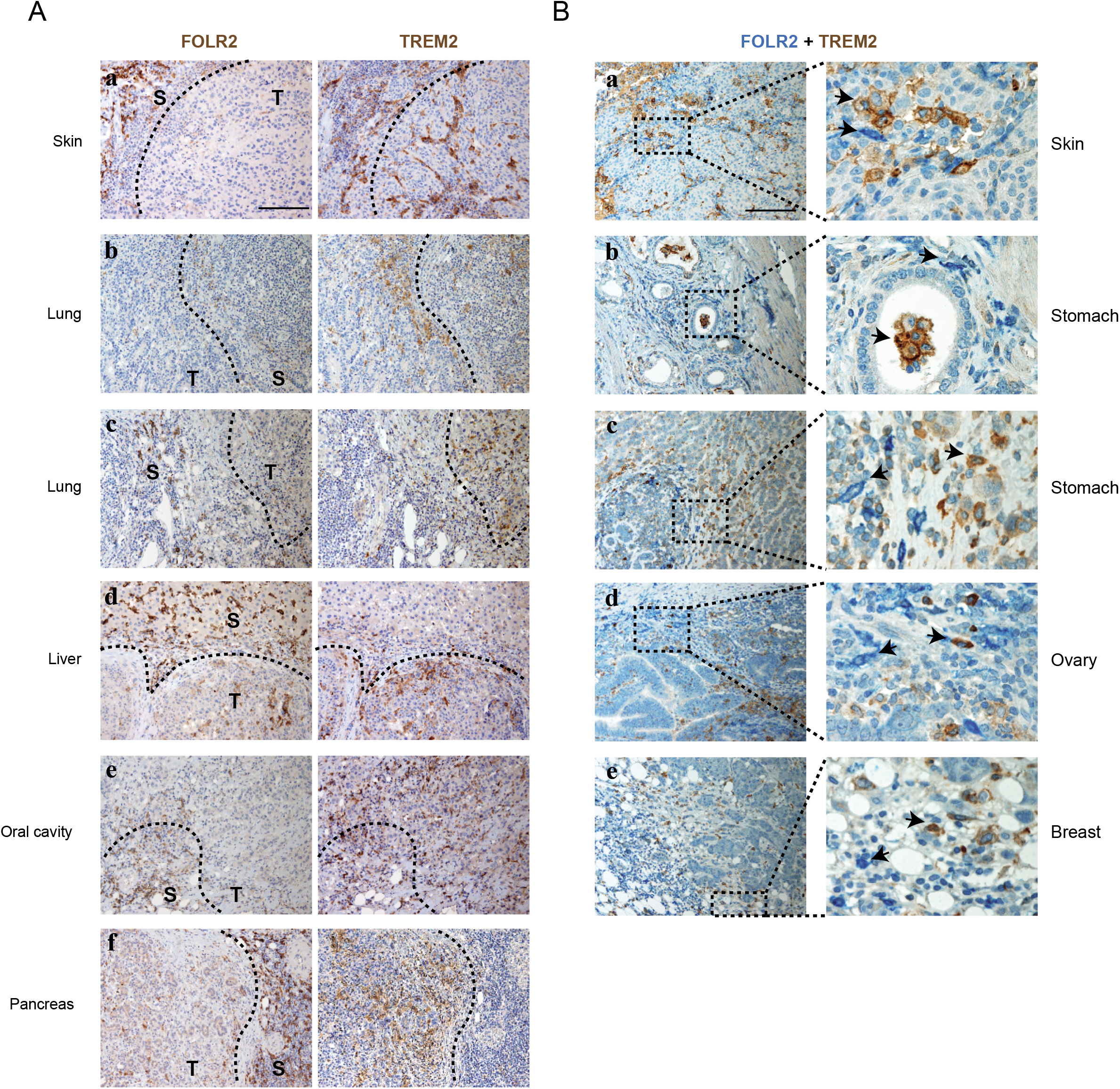
FOLR2^+^ macrophages are spatially segregated from TREM2^+^ macrophages. **A.** Spatial distribution of FOLR2^+^ TAMs and TREM2^+^ TAMs in human primary carcinomas. Serial sections are from six cases of primary cancers including one melanoma (**a**), lung carcinomas (**b, c**), hepatocellular carcinoma (**d**), oral cavity squamous cell carcinoma (**e**), pancreatic carcinoma (**f**) stained as labeled. Sections are counterstained with Mayer’s hematoxilin. Magnification= 200X, scale bar 100 micron. T = tumor and S = stroma; dotted line represents invasive margin. **B.** Sections are from cutaneous melanomas (**a**), gastric carcinomas (**b, c**), a serous ovarian carcinoma (**d**), a breast carcinoma (**e**) and stained as labeled; low (left panels) and high (right panels) magnifications are reported. Double stain from FOLR2 and TREM2 confirms a dominant mutually exclusive distribution of the two markers. Sections are counterstained with Mayer’s hematoxilin. Magnification= 100X, scale bar 200 micron (left panels); 400X, scale bar 100 micron (right panels).

### FOLR2^+^ macrophages are enriched in CD8^+^ T cells infiltrated-tumors and co-localize with lymphoid aggregates across cancers

To gain further functional insight, we next used the previously defined FOLR2 gene signature (*FOLR2/SLC40A1/SEPP1*) or *FOLR2* expression alone to correlate abundance of FOLR2^+^ macrophages with other immune and stromal cell types in the TME (Fig. 6A, Table S8). We found that the FOLR2 gene signature (or *FOLR2* expression alone) positively correlated with known players of anti-tumor immunity like CD8^+^ T cells, DCs, B cells and tertiary lymphoid structures (Fig. 6A). In contrast CADM1^+^TREM2 gene signature (*TREM2/SPP1*) or *TREM2* expression alone did not significantly correlate with T cells, CD8^+^ T cells, NK or B cells (Fig. 6A). In addition, the highest level of *FOLR2* expression in bulk tumor transcriptomes coincides, in the same tumor, with coordinated infiltration of multiple lymphocyte lineages and a gene signature for tertiary lymphoid structures (Fig. 6B). By contrast, no correlation was found when patients were stratified according to levels of *TREM2* transcript (Fig. 6B). We analyzed gene pathways represented in all genes positively correlated to *FOLR2* (or *TREM2*) expression in whole tumor transcriptome (Fig S6A). In support of our previous findings, we found a strong correlation between *FOLR2* expression in tumors and various immune pathways, including TCR signaling, antigen processing and PD-1 signaling. We also analyzed gene pathways enriched in FOLR2^+^ TAMs as compared to CADM1^+^TREM2^+^ TAMs in our bulk RNAseq dataset. We found that gene pathways enriched in FOLR2^+^ TAMs were involved in chemotaxis and functional modules of immune response regulation (Fig. S6C-D). Altogether, these results suggest that FOLR2^+^ macrophages are part of an immune contexture underlying the onset of anti-tumor immunity.

**Figure 6.**
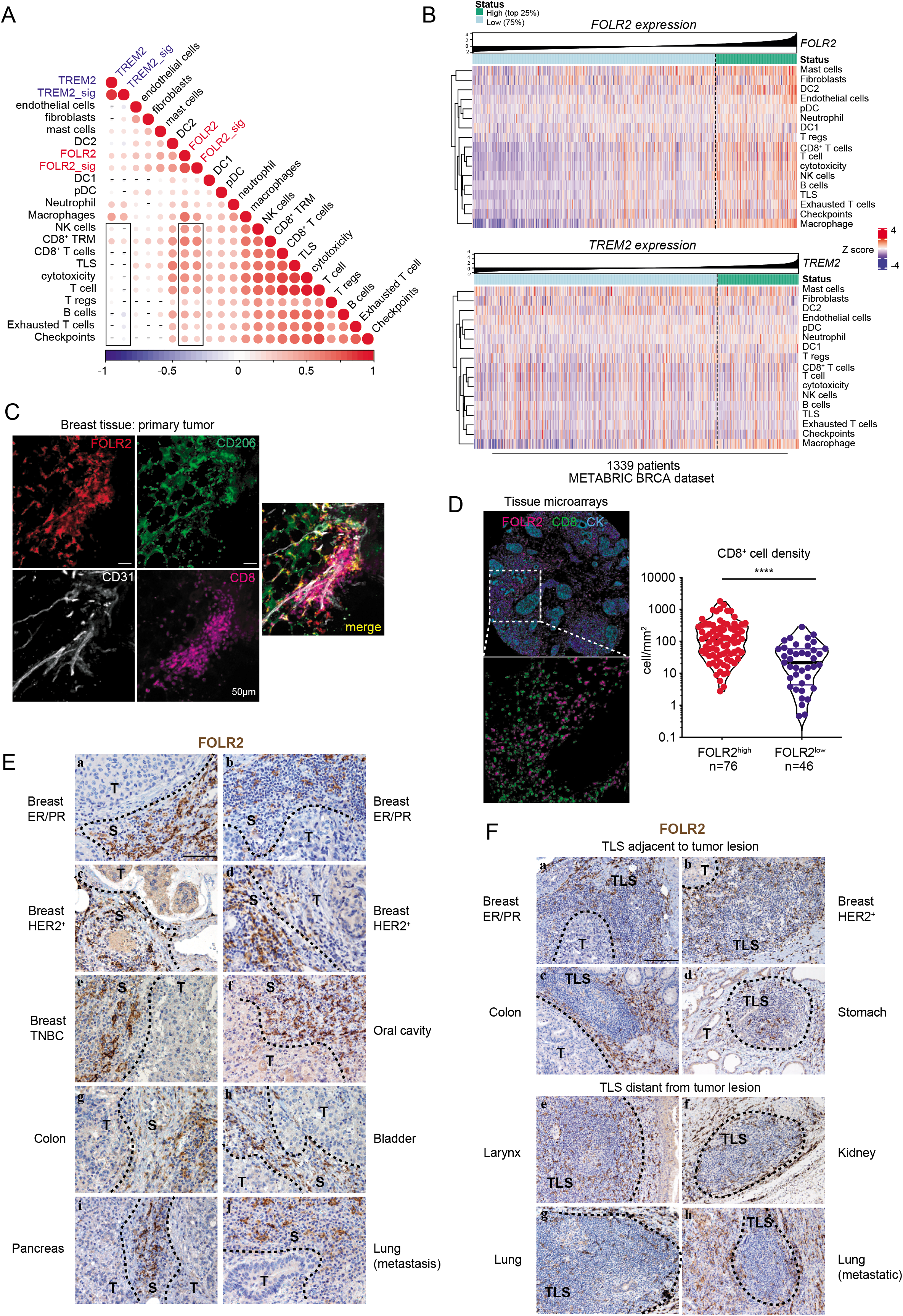
FOLR2^+^ macrophages are enriched in CD8^+^ T cells infiltrated-tumors and co-localize with lymphoid aggregates across cancers. **A-B**. Correlation map (**A**) and heatmaps (**B**) analyzing the association of FOLR2 and TREM2 genes to immune cell gene signatures in the METABRIC dataset. **C**. Representative confocal immunofluorescence images showing FOLR2^+^CD206^+^ macrophages colocalizing with CD31^+^ vessels and CD8^+^ T cell in primary breast cancer. **D**. Representative multiplex immunofluorescence images of tissue-microarray showing CD8, FOLR2 and Cytokeratin (CK) protein expression. Bottom, graph showing density of CD8^+^ T cells in tumors with high or low FOLR2^+^ cell density. **E.** Sections are from cases of primary breast carcinomas showing a luminal (**a, b**), a HER2^+^ (**c, d**) and a triple negative (**e**) phenotypes, and from cases of squamous cells carcinoma of oral cavity (**f**), colorectal carcinoma (**g**), urothelial bladder cancer (**h**), pancreatic carcinoma (**h**) and one case of metastatic carcinomas to lung (**j**), stained as labeled. Sections are counterstained with Mayer’s hematoxylin. Magnification= 200X, scale bar 100 micron (**a-l**). T = tumor and S = stroma; dotted line represents invasive margin. **F.** Sections are from primary luminal A (**a**) and HER2^+^ (**b**) breast carcinomas, colorectal carcinoma (**c**), gastric carcinoma (**d**), carcinoma of the larynx (**e**), kidney carcinoma (**f**), lung carcinoma (**g**) and lung metastatic carcinoma (**h**), stained for anti-FOLR2. FOLR2^+^ macrophages are found in tertiary lymphoid strcutures (TLS) adjacent to (**a-d**) or distant from (**e-h**) neoplastic cells. Sections are counterstained with Mayer’s hematoxilin. Magnification 100X, scale bar 200 micron (**a-h**).

Since CD8^+^ T cells have been shown to be associated to better survival in various cancer types including breast cancer (DeNardo et al., 2011)(Ali et al., 2014)(Pagès et al., 2018), we investigated whether FOLR2^+^ macrophages could be found interacting with tumor-infiltrating CD8^+^ T cells. We used confocal microscopy on tumor resection samples and found that FOLR2^+^ macrophages located near CD31^+^ vessels were closely associated with CD8^+^ T cell aggregates (Fig. 6C). To confirm the spatial association between FOLR2^+^ macrophages and CD8^+^ T cells we stained the previous tissue microarray patient cohort with both CD8 and FOLR2 and calculated their respective cellular density. FOLR2 macrophages^high^ tumor lesions had significantly higher CD8^+^ T cell density than FOLR2 macrophages^low^ lesions (Fig. 6D). Importantly, FOLR2^+^ macrophages found within peri-tumoral stroma were repeatedly enriched in lymphoid aggregates across various cancer types (Fig. 6E, S6B, Table S7) and could also be detected within tertiary lymphoid structures (Fig. 6F, S6C). Altogether, these results show that stroma-associated FOLR2^+^ macrophages are conserved across cancers and are structural component of lymphoid structures near tumor nests. These lymphoid structures are likely to be associated to ongoing immune response.

We therefore conclude that FOLR2^+^ macrophages represent a defined macrophage population associated to the onset of anti-tumoral immune responses.

## DISCUSSION

In order to understand the phenotypic and functional heterogeneity of macrophage populations in human breast cancers, we generated a single-cell atlas of myeloid cells infiltrating human luminal breast tumors. We identified two phenotypically distinct TAM subsets: TREM2^+^CADM1^+^ macrophages and FOLR2^+^ macrophages. In stark contrast with TREM2^+^ macrophage whose infiltration has been previously associated to worst clinical outcomes (Molgora et al., 2020), we show that the abundance of FOLR2^+^ TAMs is predictive of better clinical outcomes. In addition, we establish that FOLR2^+^ TAMs are spatially associated with sites of lymphocyte infiltration within the TME across cancers.

### FOLR2^+^ macrophages: a subset of TRMs associated with favorable clinical outcome

In this study, we have identified an evolutionarily conserved tissue-resident FOLR2^+^ macrophage population present in both human and mouse healthy mammary gland. In mice, breast TRMs regulate post-natal mammary gland development and remodeling (Gouon-Evans et al., 2002)(Van Nguyen and Pollard, 2002). Human FOLR2^+^ macrophages express MRC1 and LYVE1 and align to murine MRC1^+^TRMs that had been described in adult mammary gland of healthy nulliparous mice (Jäppinen et al., 2019)(Wang et al., 2020). MRC1^+^ TRMs arise from fetal precursors as demonstrated by genetic labeling at E8.5 or E13.5 in fate-mappers *Csf1r^Mer-iCre-Mer^* or *Cx3cr1*^Cre-ERT2^ mice respectively (Jäppinen et al., 2019). In addition, MRC1^+^ TRMs exhibit a self-renewing capability (Wang et al., 2020). MRC1^+^ TRMs have a non-redundant function in mammary gland development: inhibition of MRC1^+^ TRM development in *Plvap^-/-^* mice significantly impairs ductal morphogenesis during puberty (Jäppinen et al., 2019). In mice, MRC1^+^ TRMs co-exist with a minor fraction of CX3CR1^high^MRC1^-^ TRMs endowed with an intra-ductal localization (Dawson et al., 2020). Intra-ductal CX3CR1^high^MRC1^-^ macrophages develop from adult monocytes and they expand during tissue remodeling imposed by lactation (Dawson et al., 2020). Interestingly, most breast tumor-invading murine TAMs align transcriptionally to homeostatic intra-ductal CX3CR1^high^MRC1^-^ TRM (Dawson et al., 2020). In sum, these findings highlight the ontogenetic and functional diversity within mammary gland TRM subsets (Ginhoux and Guilliams, 2016).

Here we identify the FOLR2^+^ macrophages as human orthologs of murine MRC1^+^ breast TRMs. We found that human FOLR2^+^ macrophages represent the main macrophage population in healthy breast tissue. This defines FOLR2^+^ macrophages as *bona fide* mammary gland-resident macrophages. In contrast, we show that CADM1^+^TREM2^+^ macrophages are scarce in healthy tissue and increase in metastatic LN and primary tumors. We show that human CADM1^+^TREM2^+^ macrophages align with murine CX3CR1^high^MRC1^-^ TAMs. These findings are consistent with a recent study identifying TREM2^+^ macrophages in multiple cancer types (Molgora et al., 2020). The infiltration of CADM1^+^TREM2^+^ macrophages is likely to rely on influx of circulating monocytes locally attracted to tumor lesions. In support of this view, our bulk RNAseq analysis of human CADM1^+^TREM2^+^ macrophages demonstrate their transcriptional closeness to CCR2^+^ monocytes.

Do TRMs have a specific function during carcinogenesis? Recent studies have reported pro-tumorigenic activities for murine TRMs. For instance, depletion of embryonic-derived pro-fibrotic TRMs delays the progression of tumor lesions in pancreatic ductal adenocarcinoma murine models (Zhu et al., 2017). However, it is not known if this impacts on overall survival. Also, depletion of CD163^+^ TRMs in ovarian cancer reduces epithelial to mesenchymal transition and overall tumor growth (Etzerodt et al., 2020). In murine PyMT breast cancer, Franklin et al. have shown that MRC1^+^ TRMs present in healthy mammary glands, persist in developing murine breast adenocarcinoma despite dilution by incoming monocyte-derived TAMs (Franklin et al., 2014). The pro-tumorigenic function of TRMs found in pancreatic and ovarian cancers does not seem to apply to breast cancer in which MRC1^+^ TRMs are less immunosuppressive than monocyte-derived, NOTCH-dependent TAMs (Franklin et al., 2014)(Kitamura et al., 2018). MRC1^+^ TRM depletion prior to carcinogenesis did not affect tumor growth in autochthonous MMTV-PyMT or MMTV-Her2 mouse models (Franklin et al., 2014)(Linde et al., 2018) despite an effect on early cancer cell dissemination (Linde et al., 2018).

In human, the lack of specific markers distinguishing monocyte-derived TAMs versus TRMs has so far precluded an assessment of their function. To address this gap in knowledge, we have defined specific gene-signatures and surface markers enabling careful identification of macrophage subsets within tumors. To probe the association of FOLR2^+^ macrophages with clinical outcome in breast cancer we designed a gene-signature enabling us to infer FOLR2^+^ macrophages-abundance in breast cancer bulk transcriptomes. We found that the abundance of FOLR2^+^ macrophages, but not of CADM1^+^TREM2^+^ macrophages, associated with better prognosis. We provide a validation of this *in silico* finding showing that the density of FOLR2^+^ macrophages in a retrospective cohort of 122 breast cancer patients correlates with increased survival. Altogether, these results establish FOLR2^+^ macrophages as a biomarker of favorable outcome.

### Functional diversity of perivascular macrophages

We observed that FOLR2^+^ macrophages present a transcriptional signature of steady state perivascular (PV) macrophages (*LYVE1*, *MRC1*, *TIMD4*, *MAF*). Accordingly, we found that some FOLR2^+^ macrophages located in close proximity to CD31^+^ vessels. In mice, various studies have identified PV macrophages across organs including lung and skin (Chakarov et al., 2019), brain (Goldmann et al., 2016)(Sg et al., 2020), arterial wall (Lim et al., 2018), mammary gland (Jäppinen et al., 2019) and spleen (Mebius and Kraal, 2005). However, less is known about the phenotype of human PV macrophages (Lapenna et al., 2018).

In mouse models, a pro-tumoral subset of PV macrophages expressing the angiopoietin receptor TIE2 has been implicated in angiogenesis and tumor spreading (Lewis et al., 2016)(De Palma et al., 2005)(Pucci et al., 2009). Mechanistically, TIE2^+^ PV macrophages release VEGFA that favors tumor cell intravasation and metastasis by reducing tight-junctions in tumor blood vessels promoting permeabilization of the vascular wall (Harney et al., 2015). TIE2^+^ PV macrophages respond to endothelial-derived angiopoietin 2 (ANG2) engaging the TIE2 receptor thereby supporting PV positioning and pro-angiogenic function (Mazzieri et al., 2011). It is unclear if FOLR2^+^ PV macrophages described in this study align to TIE2^+^ PV macrophages. We do not favor this hypothesis for two reasons. First, we were not able to document TIE2 expression in single cell- and bulk-RNAseq analysis of FOLR2^+^ macrophages. Second, TIE2^+^ PV macrophages ontogeny relies on the progressive infiltration of a specialized subset of pro-angiogenic TIE2^+^ monocytes (De Palma et al., 2005)(Pucci et al., 2009)(Coffelt et al., 2010)(Arwert et al., 2018). This contrasts with our data evidencing FOLR2^+^ macrophages are TRMs. Therefore, further experimental efforts are needed to disentangle the heterogeneity of PV macrophages.

### A role for FOLR2^+^ macrophages in promoting lymphocyte infiltration and anti-tumor immunity?

A recent study has identified a subset of FOLR2^+^ TRMs associated to human hepatocellular carcinoma and resembling fetal-liver macrophages (Sharma et al., 2020). CD86 expression in hepatic FOLR2^+^ macrophages suggest that they might be interacting with CTLA4^+^ regulatory T cells. To probe the function of FOLR2^+^ macrophages in breast cancer, we used our gene signature to infer FOLR2^+^ macrophages-abundance in bulk tumor microarray and correlate it to the infiltration of other immune cells. We found that the infiltration of FOLR2^+^ macrophages, in contrast with TREM2^+^ macrophages infiltration, positively correlated with tumor-infiltrating lymphocytes including CD8^+^ T cell, B cells as well as DCs. In addition, we found by confocal imaging that FOLR2^+^ macrophages co-localized with CD8^+^ T cell aggregates in the vicinity of endothelial cells. We validate the correlation between FOLR2 and CD8^+^ T cell abundance by multispectral imaging: tumor lesions highly infiltrated with FOLR2^+^ macrophages had significantly higher CD8^+^ T cell-density. Since CD8^+^ T cell infiltration correlates with better survival probability in many cancers including breast cancer (DeNardo et al., 2011)(Ali et al., 2014), these results suggest that FOLR2^+^ macrophages participate to the onset of anti-tumor immunity.

FOLR2^+^ macrophages have been described in human tissues including fetal liver, placenta, colon (Samaniego et al., 2014)(Sharma et al., 2020)(Thomas et al., 2021). Here we extend these observations in healthy mammary gland and breast cancer and multiple other cancer types. Importantly, the association of FOLR2^+^ macrophages with peri-tumoral, stromal lymphoid aggregates is found in multiple cancer types. In some instances, we report FOLR2^+^ macrophages infiltrating organized tertiary lymphoid structures comprising B and T cells. Murine PV macrophages have been described as possible regulators of lymphocyte infiltration during inflammation (Natsuaki et al., 2014) and auto-immunity (Mohan et al., 2017). It can be hypothesized that FOLR2^+^ macrophages could regulate the infiltration of CD8^+^ T cells by different mechanisms: directly by delivering chemokines attracting T cells (Dangaj et al., 2019), or indirectly by delivering inflammatory cytokines to endothelial cells or growth factor to pericytes (Minutti et al., 2019). Further studies are needed to identify the mechanisms by which FOLR2^+^ macrophages regulate lymphocyte infiltration in tumors, a key event for the development of efficient anti-tumor immune responses.

## ACKNOWLEDGMENTS

We thank the Institut Curie Cytometry platform for technical help. We are grateful to all members of the TransImm team for helpful discussions and comments. We also thank Ana-Maria Lennon-Dumenil and Federica Benvenuti for helpful and precious advices. We thank Charlotte Martinat for her help in handling patient samples. TransImm team is supported by the SiRIC-Curie Program (grant INCa-DGOS-12554), the LabEx DCBIOL (ANR-10-IDEX-0001-02 PSL and ANR-11-LABX-0043), and the Center of Clinical Investigation (CIC IGR-Curie 1428). High-throughput sequencing was performed by the ICGex NGS platform of the Institut Curie supported by the grants ANR-10-EQPX-03 (Equipex) and ANR-10-INBS-09-08 (France Génomique Consortium) from the Agence Nationale de la Recherche (“Investissements d’Avenir” program), by the ITMO-Cancer Aviesan (Plan Cancer III) and by the SiRIC-Curie program (SiRIC Grant INCa-DGOS-4654). W.R. is supported by the grant IG23179, Associazione Italiana per la Ricerca sul Cancro.

## AUTHOR CONTRIBUTIONS

Conceptualization, R.N.R., E.P. and J.H; Methodology, R.N.R., Y.K.M, Y.G.F.; Formal Analysis, W.R., Y.K.M., C.B.; Investigation, R.N.R., Y.K.M, Y.G.F., N.G.N., J.B.T., C.B., M.B., J.D., P.C., F.K., L.L.N., C.A.D., F.G.,L.V., E.D., B.B., J.G., S.Z., W.V.; Resources, S. B., S.L., M.B., C.S., S.V., L.S., A.N., D.M., A.V.S, F.R; Writing – Original Draft, J.H.; Writing – Review & Editing, J.H., P.G., S.Z., E.P., R.N.R. E.D.; Visualization, J.H., R.N.R. and Y.K.M. Supervision, J.H.; Funding Acquisition, J.H., E.P.

## EXPERIMENTAL MODEL AND SUBJECT DETAILS

### Human tumor samples

Primary tumors, juxta-tumor tissues, metastatic and non-metastatic tumor-draining LNs were surgically resected from luminal breast cancer patients at the Institut Curie Hospital (Paris, France), in accordance with institutional ethical guidelines and informed consent was obtained (approved by the Ethical Committee of Curie Institute, CRI-0804-2015). Patients clinical and pathologic information are summarized in Table S1. Lymph node metastases were assigned by the pathology department of Institut Curie and further confirmed by using pan-cancer/CD45 immune detection by flow cytometry. FOLR2 and TREM2 Immunohistochemistry was performed on tumor sections retrieved from the archive of the Pathology Unit, ASST Spedali Civili di Brescia. Patients clinical and pathologic information are summarized in Table S7.

### Mice

Transgenic PyMT mice (MMTV-PyMT^634Mul^)(Davie et al., 2007) were maintained on C57Bl/6 background and were bred and maintained in specific pathogen-free in Curie Institute animal facility in accordance with Curie Institute guidelines. Healthy C57BL/6J female mice were obtained from Charles River Laboratories, maintained in a non-barrier facility and included at 8-12 weeks of age for experimental procedures. Animal care and use for this study were performed in accordance with the recommendations of the European Community for the care and use of laboratory animals (2010/63/UE). Experimental procedures were specifically approved by the Ministère de l’Enseignement Supérieur et de la Recherche (authorization number 2016-06.150) in compliance with the international guidelines.

## METHOD DETAILS

### Human samples processing

Patient samples were processed as previously described (Núñez et al., 2020). In brief, freshly resected human samples were cut into small fragments and digested with 0.1 mg/ml Liberase TL (Roche) and 0.1 mg/ml DNase (Roche) in CO2-independent medium (GIBCO) + 0,4 g/l of human albumin (Vialebex) for 30 min at 37°C. Single cell suspension of dissociated tissues were filtered on a 40-μm cell cell strainer (BD Biosciences), washed with CO2-independent medium + 0.4 g/l of human albumin and resuspended in medium for cell counting estimation.

### Flow cytometry analysis

After tissue processing and cell counting, cell suspensions were stained with Live/Dead Fixable Aqua Dead Cell Stain Kit (Life Technologies) in 1X phosphate-buffered saline (PBS) for 10 minutes at room temperature. After incubation cells were washed in cold PBS + 0.5% bovine serum albumin (BSA) + 2mM EDTA and stained with fluorescent antibodies (Table S9) in the presence of Fc-receptors blocking reagent (Miltenyi) during 30 minutes at 4°C. Cell suspensions were subsequently washed and submitted to intracellular staining using fixation/permeabilization kit (eBiosciences/Thermo Fischer) accordingly to the manufacturer’s instructions. Data acquisition was performed using an LSR-Fortessa (BD), compensation and analysis were done using FlowJo software (TreeStar).

### Mass cytometry staining and data analysis

For mass cytometry, pre-conjugated or purified antibodies were obtained from Invitrogen, Fluidigm (pre-conjugated antibodies), Biolegend, eBioscience, Becton Dickinson or R&D Systems as listed in Table S9. For some markers, fluorophore-conjugated or biotin-conjugated antibodies were used as primary antibodies, followed by secondary labeling with anti-fluorophore metal-conjugated antibodies (such as the anti-FITC clone FIT 22) or metal-conjugated streptavidin, produced as previously described (Becher et al., 2014). Briefly, patient lymph nodes cell suspension (around 30×10^6^ cells/well in a U-bottom 96 well plate; BD Falcon, Cat# 3077) were washed once with 200 mL FACS buffer (4% FBS, 2mM EDTA, 0.05% Azide in 1X PBS), then stained with 100 mL 200 mM cisplatin (SigmaAldrich, Cat# 479306-1G) for 5 min on ice to exclude dead cells. Cells were then washed with FACS buffer and once with PBS before fixing with 200 mL 2% paraformaldehyde (PFA; Electron Microscopy Sciences, Cat# 15710) in PBS overnight or longer. Following fixation, the cells were pelleted and resuspended in 200μL 1X permeabilization buffer (Biolegend, Cat# 421002) for 5 min at room temperature to enable intracellular labeling. Bromoacetamidobenzyl-EDTA (BABE)-linked metal barcodes were prepared by dissolving BABE (Dojindo, Cat# B437) in 100mM HEPES buffer (GIBCO, Cat# 15630) to a final concentration of 2 mM. Isotopically-purified PdCl2 (Trace Sciences) was then added to the 2 mM BABE solution to a final concentration of 0.5 mM. Similarly, DOTA-maleimide (DM)-linked metal barcodes were prepared by dissolving DM (Macrocyclics, Cat# B-272) in L buffer (MAXPAR, Cat# PN00008) to a final concentration of 1 mM. RhCl3 (Sigma) and isotopically-purified LnCl3 was then added to the DM solution at a final concentration of 0.5 mM. Six metal barcodes were used: BABE-Pd-102, BABE-Pd-104, BABE-Pd-106, BABE-Pd-108, BABE-Pd-110 and DMLn-113. All BABE and DM-metal solution mixtures were immediately snap-frozen in liquid nitrogen and stored at 80C. A unique dual combination of barcodes was chosen to stain each tissue sample. Barcode Pd-102 was used at a 1:4000 dilution, Pd-104 at a 1:2000, Pd-106 and Pd-108 at a 1:1000, and Pd-110 and Ln-113 at a 1:500. Cells were incubated with 100 mL barcode in PBS for 30 min on ice, washed in permeabilization buffer and then incubated in FACS buffer for 10 min on ice. Cells were then pelleted and resuspended in 100 mL nucleic acid Ir-Intercalator (MAXPAR, Cat# 201192B) in 2% PFA/PBS (1:2000), at room temperature. After 20 min, cells were washed twice with FACS buffer and twice with water before being resuspended in water. In each set, the cells were pooled from all tissue types, counted, and diluted to 0.5×106 cells/mL. EQ Four Element Calibration Beads (DVS Science, Fluidigm) were added at a 1% concentration prior to acquisition. Cell data were acquired and analyzed using a CyTOF Mass cytometer (Fluidigm). The CyTOF data were exported in a conventional flow-cytometry file (.fcs) format and normalized using previously-described software (Finck et al., 2013). Events with zero values were randomly assigned a value between 0 and −1 using a custom R script employed in a previous version of the mass cytometry software (Newell et al., 2012). Cells for each barcode were deconvolved using the Boolean gating algorithm within FlowJo. The CD45^+^Lin(CD3/CD19/CD20)^-^HLA-DR^+^ population of PBMC were gated using FlowJo and exported as a .fcs file.

### Human and mouse sample processing for single cell sequencing

#### Mouse tumors

from PyMT or wild-type littermate control were cut in small pieces and digested for 1h at 37°C in CO2-independent medium (Gibco) containing 150 μg/mL DNAse and 75 μg/mL Liberase TL (Roche). Cell suspensions were then obtained by filtering on a 40μM cell-strainer (BD). Red blood cells were subsequently lysed with red blood lysis buffer. Finally, cells were resuspended in cold PBS containing 2mM EDTA (Invitrogen) and 5% bovine serum albumin (BSA; Sigma) for further assays.

#### Human tumors

were processed according to previously published method (Leruste et al., 2019)(Bourdely et al., 2020). Briefly, after tissue processing, dissociation and cell counting, cell suspensions were maintained on ice and stained for FACS-sorting with antibodies depicted on Table S9 and DAPI. Cells were isolated using FACS-ARIA III (BD) cell sorter using gating strategies shown at Fig. S2C and collected in cold PBS+0,04% of BSA for cell counting. PBMC were obtained from fresh blood samples by density gradient centrifugation using Lymphoprep (Stemcell Technologies) according to the manufacturer instructions, then washed and resuspended in CO2-independent medium + 0.4 g/l of human albumin prior FACS-sorting. Blood monocytes were also collected in cold PBS+0.04% of BSA before cell counting. All tissues were processed within 1 hour after tumor resection, and sorted cells were loaded in a 10x Chromium chip instrument within 6 hours.

### Single-cell RNA-sequencing

Human and mouse cellular suspensions (3000 to 8000 cells) were loaded on a 10x Chromium Controller (10X Genomics) according to manufacturer’s protocol based on the 10x GEMCode proprietary technology. Single-cell RNA-Seq libraries were prepared using Chromium Single Cell 3’ v2 or v3 Reagent Kit (10x Genomics) according to manufacturer’s protocol. Briefly, the initial step consisted in performing an emulsion where individual cells were isolated into droplets together with gel beads coated with unique primers bearing 10X cell barcodes, unique molecular identifiers (UMI), and poly(dT) sequences. Reverse transcription reactions were engaged to generate barcoded full-length cDNA followed by the disruption of emulsions using the recovery agent and cDNA clean up with DynaBeads MyOne Silane. Bulk cDNA was amplified using a GeneAmp PCR System 9700 with 96-Well Gold Sample Block Module (Applied Biosystems) (98 °C for 3 min; cycled 11/12 ×: 98 °C for 15 s, 63 °C for 20 s and 72 °C for 1 min; held at 4 °C). Amplified cDNA product was cleaned up with the SPRI select Reagent Kit (Beckman Coulter). Indexed sequencing libraries were constructed using the reagents from the Chromium Single Cell 3’ v3 Reagent Kit, following these steps: (1) fragmentation, end repair, and a-tailing; (2) size selection with SPRI select; (3) adaptor ligation; (4) post ligation cleanup with SPRI select; (5) sample index PCR and cleanup with SPRI select beads. Library quantification and quality assessment was performed using Qubit fluorometric assay (Invitrogen) with dsDNA HS (High Sensitivity) Assay Kit and Bioanalyzer Agilent 2100 using a High Sensitivity DNA chip (Agilent). Indexed libraries were pooled according to number of cells and sequenced on a NovaSeq 6000 (Illumina) using paired-end 28 × 91 bp. A depth around 50,000 reads per cell was obtained.

### Single-cell RNA-sequencing data processing and analysis

#### Alignment and Raw Expression Matrix Construction

Human and mouse raw sequencing data were respectively aligned on reference genome GRCh38/84 and GRCm38/84 (genome assembly/ENSEMBL release) using 10X software CellRanger (Version 3.0.2) with default parameters. Gene expression counts for individual cells were generated using cellranger count. <u>Cell Selection, Filtering and Normalization.</u> For both human and mouse raw count matrix, we kept cells expressing at least 200 genes. Cells with mitochondrial content greater than 10 % were removed. After this quality control, data were normalized by total counts following the Seurat 3 R pipeline. Cells identified as contaminant of the gating strategy or with high heat-shock-protein or ribosomal coding genes content were filtered out. <u>Variable Gene Selection and Sample Merging.</u> For sample merging we applied a VST (Variance Stabilizing Transformation) method selecting for the most variable genes. We performed the Seurat V3 integration pipeline using the most 8000 genes for the ten human samples. The 3000 most variable genes were used to merge the two mice samples. For both anchors selection and integration steps we used the default parameters of Seurat V3 functions. <u>Dimensionality reduction and Visualization.</u> Data were scaled by applying a regression using as variation factors, the total UMI counts, the percent of expressed mitochondrial genes, the origin sample and tissue of each cell and the version of CellRanger chemistry kit used for sequencing. Heatmaps are showing z-scores of this scaled matrix. The UMAP visualization was built using respectively the 50 and 30 most informative components of the PCA for Human and Mouse. <u>Clustering and Differential Gene Expression Analysis.</u> The clustering was processed by constructing a Shared Nearest Neighbor (SNN) Graph. The 20 neighbors of each cell were first determined. The resulting KNN graph was used to construct the SNN graph by calculating the neighborhood overlap (Jaccard index) between every cell and its 20 nearest neighbors. Clustering was then applied on this graph using the Louvain graph-based algorithm. Differential gene expression analysis was applied on each sample log normalized matrix. We used the Seurat function *FindAllMarkers()*, with a Logistic Regression test, and adding as variation factors, the origin sample and tissue of each cell and the version of CellRanger sequencing kit used. Only genes expressed in more than 10% of the cells in a cluster and having at least 0.10 of log Fold-Change between compared groups were tested. We were able to detect low signals produced by genes with dropouts. For the volcano plots only the first condition was kept. At the end, only genes with a significative adjusted p-value (pv<0.05, false discovery rate (FDR) adjusted p-value) were kept and used to define each cluster.

#### Human-Mouse Merging and Seurat Label Transfer Prediction Score

To compare mouse and human macrophages, we selected conserved orthologue genes between both species. Corresponding gene symbols table was provided by the Mouse Genome Informatics database (Table S10). For genes with more than one corresponding orthologue in the other species, we took the mean expression of all orthologues. Seurat label transfer scoring algorithm was applied from mouse (as query) to human dataset (as reference). For the anchors searching step, we used the 10 000 most variable genes of the reference dataset. The following steps of the Seurat pipeline was applied with default parameters.

#### Processing of published dataset

We downloaded the dataset of *Azizi et al, Cell 2018* (GSE114725). We used the classical pipeline for single cell analysis of Seurat V3 (without integration correction) from the raw count matrix (supplementary file GSE114725_rna_raw.csv.gz). We next perform Louvain graphbased clustering. At the resolution 0.9 we obtained 39 clusters: 14 clusters of T cells, 6 clusters of B cells, 5 clusters of NK cells, 1 cluster of pDCs, DC1, DC2, CD16^+^ monocytes, CD14^+^ monocytes, neutrophils or mast cells, 3 clusters of macrophages and 4 clusters of contaminating cells. We merged clusters of the same immune cell types, except for the macrophages clusters and removed contaminating clusters (Fig. S3D). We downloaded the dataset of *Han et al*, Cell 2018 (GSE108097). We integrated 2 samples of virgin mammary gland and 1 sample of pregnant mammary gland from the raw data (supplementary file GSE108097_RAW.tar). We used the same pipeline described above including the integration step. Fig. S3C shows clustering at the resolution 0.1.

### Bulk RNA-sequencing

After tissue processing and cell count, cell suspensions were washed in cold PBS + 0.5% BSA + 2mM EDTA and submitted to surface antibodies staining (Table S9) in the presence of Fc-receptors blocking (FcR Blocking Reagent; Miltenyi) for 30 minutes at 4°C. Myeloid cell subsets were isolated using FACS-ARIA III (BD) cell sorter considering gating strategies shown at figure S2A and directly collected on lysing TCL buffer (QIAGEN) containing 1% of beta-mercaptoethanol before storage at −80°C. RNA were extracted and isolated using the Single Cell RNA purification kit (Norgen, Cat#51800) according to the manufacturer’s instructions. After extraction, total RNA was analyzed using Agilent RNA 6000 Pico Kit on the Agilent 2100 Bioanalyzer System. RNA quality was estimated based on capillary electrophoresis profiles using the RNA Integrity Number (RIN). RNA sequencing libraries were prepared using the SMARTer Stranded Total RNA-Seq Kit v2 - Pico Input Mammalian (Clontech/Takara). The input quantity of total RNA was comprised between 1 and 22ng. This protocol includes a first step of RNA fragmentation, using a proprietary fragmentation mix at 94°C. The time of incubation was set up for each sample, based on the RNA quality, and according to the manufacturer’s recommendations. After fragmentation, indexed cDNA synthesis was performed. Then the ribodepletion step was performed, using probes specific to mammalian rRNA. PCR amplification was finally achieved to amplify the indexed cDNA libraries, with a number of cycles set up according to the input quantity of tRNA. Library quantification and quality assessment was performed using Qubit fluorometric assay (Invitrogen) with dsDNA HS (High Sensitivity) Assay Kit and LabChip GX Touch using a High Sensitivity DNA chip (Perkin Elmer). Libraries were then equimolarly pooled and quantified by qPCR using the KAPA library quantification kit (Roche). Sequencing was carried out on the NovaSeq 6000 (Illumina), targeting between 10 and 15M reads per sample and using paired-end 2 x 100 bp.

### Bulk RNA-sequencing data processing and analysis

The raw sequencing data was initially aligned on the human reference genome hg19, using STAR aligner (v2.5.3a) (Dobin et al., 2013). Raw read counts matrix made also with STAR (using the parameter --quantMode GeneCounts). FastQ files quality control were applied with FastQC (removing of adapters and low-quality bases). Non-expressed genes (the sum of counts in all samples less than 2) and lowly expressed genes (background; log2 of the average of raw counts in all samples less than 2) were removed from the raw read count matrix. For differential expression analysis, we used the R package DESeq2 (version 1.24.0)(Love et al., 2014) with a p-value correction. The DESeq matrix was designed using the information of the patient and the cell type (following the formula design = ~ batch + condition). For the normalization, we used the median of ratios method (Anders and Huber, 2010) and the rlog transformation for visualization and clustering as proposed in the DESeq2 tutorial (Love et al., 2014).

### Immunohistochemistry

Paraffin-embedded tissue blocks were cut with a microtome into fine slivers of 3 microns. Immunohistochemistry was processed in a Bond RX automated (Leica) with Bond Polymer refine detection kit (Leica, DS9800). Antigen retrieval was performed in BOND Epitope Retrieval Solution 1 (Leica, AR9961). Primary antibody APOE (Abcam; ab52607) was incubated 30 minutes at room temperature. Slides were counterstained with hematoxylin before mounting with resin. Images were acquired by using Digital Pathology slide scanner (Ultra Fast Scanner 1.8, Philips)

FOLR2 expression was tested on human tissues by using immunohistochemistry. Sample included reactive lymph nodes (n=7), primary carcinomas (n=47), metastatic tumor draining lymph nodes (n=7) and distant metastasis to lung (n=8) and liver (n=11) from different primary site (breast, bladder, gastrointestinal, lung, kidney and skin) (Table S7) retrieved from the archive of the Pathology Unit, ASST Spedali Civili di Brescia. Briefly, anti-FOLR2 (clone OTI4G6, 1:100, ThermoFisher SCIENTIFIC) and anti-TREM2 (clone D814C, 1:100, Cell Signaling Technology) antibodies were revealed using Novolink Polymer (Leica) followed by DAB. For double staining, FOLR2 was combined with anti-CD3 (clone LN10, 1:70, Leica Biosystem), anti-CD20 (clone L26, 1:200, Leica Biosystem), anti-CD31 (clone PECAM-1, 1:50, Leica) and anti-TREM2. Briefly, after completing the first immune reaction, the second immune reaction was visualized using Mach 4 MR-AP (Biocare Medical), followed by Ferangi Blue. Localization of FOLR2^+^ cells within tertiary lymphoid structures (TLS) was confirmed by double for the B-cell marker CD20 and the T-cell marker CD3.

### Immunofluorescence on fixed tumor slices

Biopsies were fixed overnight at 4°C in a Periodate-Lysine-Paraformaldehyde solution (0.05 M phosphate buffer containing 0.1 M L-lysine [pH 7.4], 2 mg/ml NaIO4, and 10 mg/ml paraformaldehyde). Fixed tumors were then embedded in 5% low-gelling-temperature agarose (type VII-A, Sigma-Aldrich) and cut into 400 μm-thick slices as previously described (Peranzoni et al., 2018). Tumor slices were stained for 15 minutes at 37°C with antibodies shown at Table S9. All antibodies were diluted in PBS and used at a concentration of 5 μg/ml, except anti-FOLR2 and CD31 antibodies that were used at 10 μg/ml. Z-stack images of 5×5 fields were taken with a 10x water immersion objective (10x/0.3 N.A.) on an inverted spinning disk confocal microscope (IXplore, Olympus). Virtual slices were reconstituted and analyzed with the ImageJ software.

### Multispectral immunofluorescence on paraffin-embedded tissues

Paraffin-embedded tissue blocks were cut with a microtome into fine slivers of 5 microns. Immunostaining was processed in a Bond RX automated (Leica) with Opal™ 7-Color IHC Kits (Akoya Biosciences, NEL821001KT) according to the manufacturer’s instructions using antibodies shown at Table S9. Tissue sections were coverslipped with Prolong™ Diamond Antifade Mountant (ThermoFisher) and stored at 4°C. Subsequently, slides were scanned using the Vectra^®^ 3 automated quantitative pathology imaging system (Vectra 3.0.5; Akoya Biosciences). Multispectral images were unmixed and analysed using the inForm Advanced Image Analysis Software (inForm 2.4.6; Akoya Biosciences)

### Tissue Microarray (TMA) multispectral immunofluorescence

Paraffin-embedded tissue microarray from breast cancer patients (n=122 spots) were obtained commercially (AMSBIO, England). Immunostaining was processed in a Bond RX automated (Leica) with Opal™ 7-Color IHC Kits (Akoya Biosciences, NEL821001KT) according to the manufacturer’s instructions (Table S9). After immunostaining, slides were submitted to DAPI staining, washed and coverslipped with Prolong™ Diamond Antifade Mountant (ThermoFisher). Subsequently, slides were scanned using the Vectra^®^ 3 automated quantitative pathology imaging system (Vectra 3.0.5; Akoya Biosciences). Multispectral images were unmixed using the inForm Advanced Image Analysis Software (inForm 2.4.6; Akoya Biosciences) and analyzed by HALO software for immune subsets quantification.

### METABRIC data analysis

METABRIC (METABRIC Group et al., 2012) gene expression data, as well as clinical and sample level metadata were downloaded from cBioPortal. We annotated patient breast cancer subtype by defining TNBCs as those with a negative ER and HER2 status. HER2 positive patients were defined as any patients that had a HER2 positive status variable. ER/PR positive patients were defined as being HER2 negative but either ER or PR status positive. TNBCs with a positive PR IHC status were removed. Patients that died of other causes not related to their disease, as well as patients with breast sarcomas were removed. TNBC expression data was submitted to the TNBC type (Chen et al., 2012) algorithm that removed a further 6 patients from the TNBC cohort (MB-3297, MB-7269, MB-5008, MB-6052, MB-0179, MB-2993. The final cohort consisted of 1339 samples (168 TNBC, 204 HER2 and 967 ER/PR). To score individual samples for gene signatures of interest, we Z-score normalised the gene expression data, and then calculated a mean level of expression across signature genes. MCPcounter (1.2.0) was used to infer the abundance of immune and stromal cell populations in each sample. To stratify patients according to gene signature scores, we used a 75% cut-off therefore defining 25%of patients as “high” scorers. We carried out survival analysis using the survival (3.1-12) and survminer (0.4.6) packages in R. We computed pairwise Spearman correlations between signatures of interest and plotted these as a correlation heatmap using the corrplot R package (0.84). Correlations that were statistically non-significant (p<0.05) were marked with a dash (-). MCP counter-estimated immune cell abundance. Comparing MCP counter cell types between patients stratified into high and low groups by a particular signature, we used Wilcoxon tests with P values adjusted using the Benjamini-Hochberg method. We used ggplot2 (3.3.0), pheatmap (1.0.12) and corrplot (0.84) packages for plotting data.

### CPTAC data analysis

Data used in this publication were generated by the Clinical Proteomic Tumor Analysis Consortium (NCI/NIH). Log-ratio normalised proteomic data including clinical information and RNA sequencing data from the CPTAC BRCA study were downloaded (http://linkedomics.org/data_download/TCGA-BRCA/). BRCA patient samples annotated as either ER or PR positive, and HER2 negative, were selected for downstream analysis. Patients were then stratified according to a 25% cut-off by FOLR2 protein expression and univariate Cox regression analysis was carried out. Logrank test scores were plotted on Kaplan-Meier plots.

### GTEx to TCGA comparison

To compare TREM2 and FOLR2 mRNA expression between non-disease healthy tissue, tumour-adjacent normal tissue, and tumour tissue, we harnessed datasets from GTEx and TCGA consortia. For breast, colon and lung studies, we downloaded normalised transcriptomic data from Github (https://github.com/mskcc/RNAseqDB/tree/master/data/normalized). We used non-parametric, Wilcoxon T tests to compare TREM2 and FOLR2 expression between tissue types. **The Genotype-Tissue Expression** (GTEx) Project was supported by the <u>Common Fund</u> of the Office of the Director of the National Institutes of Health, and by NCI, NHGRI, NHLBI, NIDA, NIMH, and NINDS.

### Statistical analysis

The tests used for statistical analyses are described in the legends of each concerned figure and have been performed using GraphPad Prism v8 or R v3.4. Symbols for significance: ns, non-significant; *, <0.05, **, <0.01; ***, <0.001; ****, <0.0001. For each experimental group, n represent the number of subjects within each group.

## Supplemental figure and table legends

**Figure S1.**
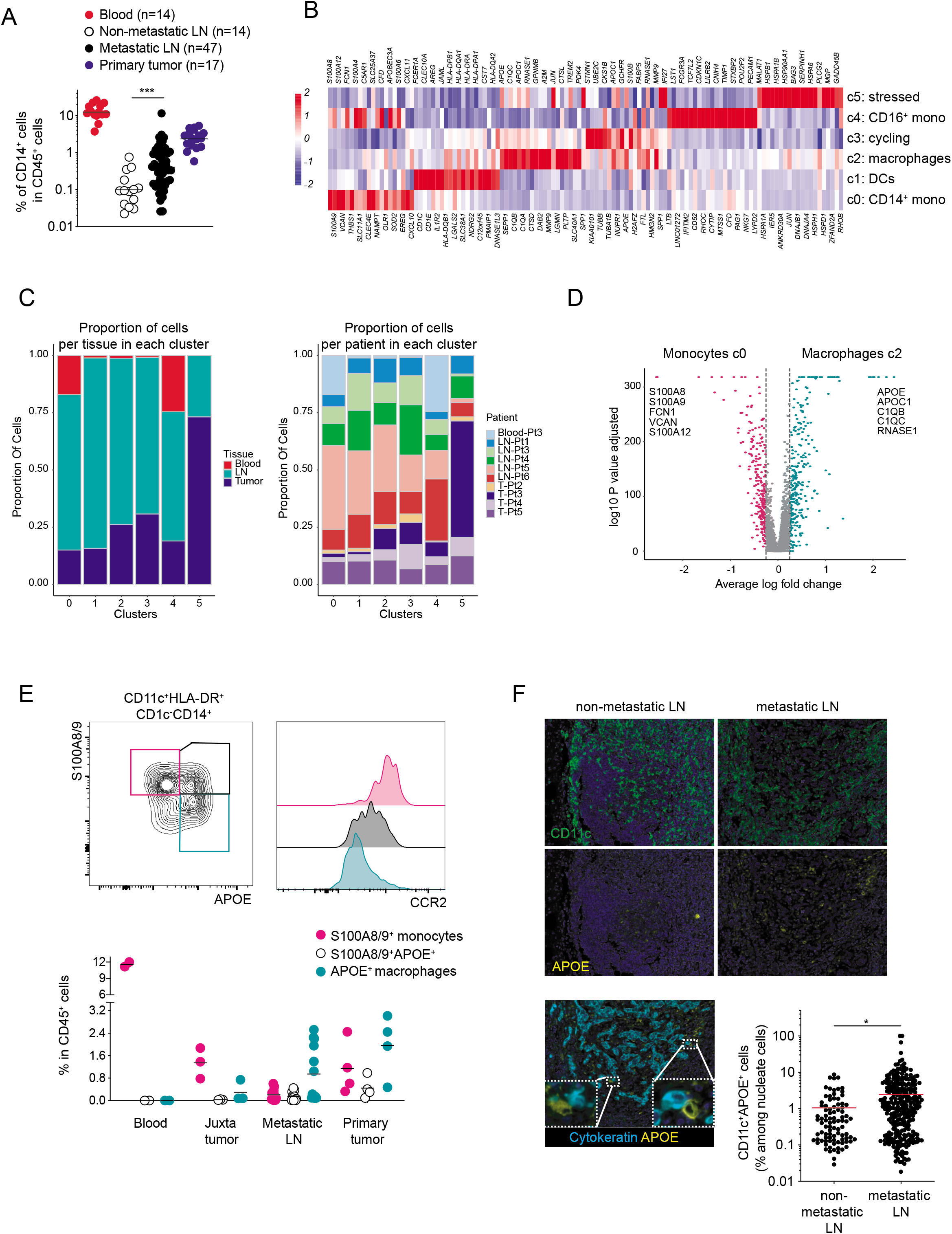
**A.** Flow cytometry quantification of CD14^+^ cells among CD45^+^ cells in blood, non-metastatic lymph nodes, metastatic lymph nodes and primary tumors of treatment-naïve luminal breast cancer patients. **B.** Heatmap showing normalized expression of the top 10 most variable genes for each myeloid cell cluster defined in Figure 1D. **C.** Proportion of cells per tissue or per patients in each cluster. **D.** Volcano plot showing D.E.G. between clusters 0 and 2. Selected genes among the Top25 were depicted. **E.** Representative contour plot showing the expression of S100A8/9, APOE and CCR2 in CD11c^+^HLA-DR^+^CD1c^-^CD14^+^ cells measured by flow cytometry. Quantification of each gates in blood, juxta-tumor tissue, metastatic lymph nodes and primary tumors of treatment-naïve luminal breast cancer patients. **F.** Immunofluorescence images of APOE^+^ macrophages and cytokeratin^+^ tumor cells in non-metastatic and metastatic lymph nodes. Quantification of CD11c^+^APOE^+^ cells among nucleated cells from multispectral images of non-metastatic and metastatic lymph nodes.

**Figure S2.**
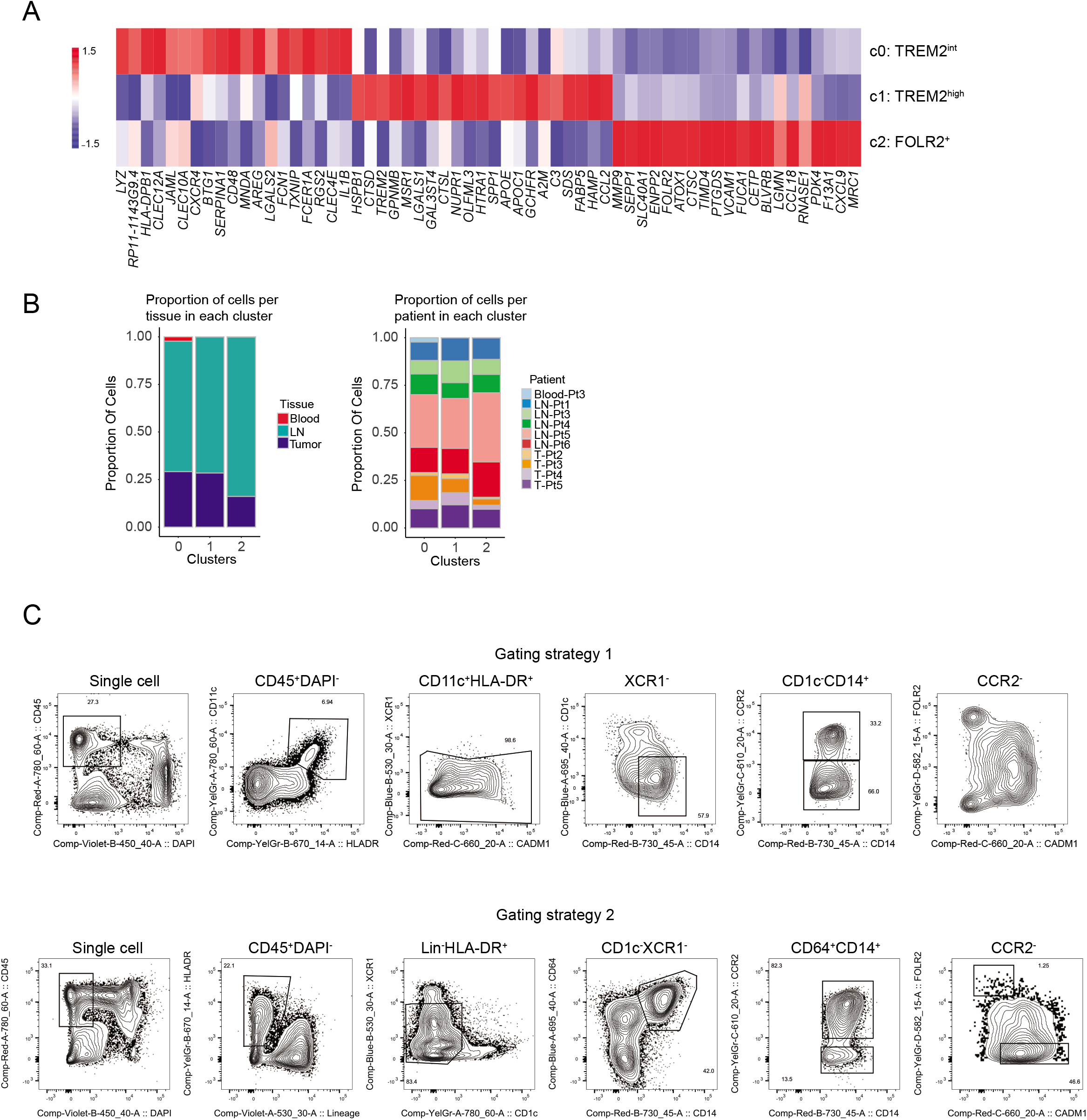
**A.** Heatmap showing normalized expression of the top 20 most variable genes for each macrophage cluster defined in Figure 2A. **B.** Proportion of cells per tissue or per patients in each cluster. **C.** Gating strategies for FACS-sorting of FOLR2^+^ and CADM1^+^ macrophages isolated from metastatic lymph nodes or primary tumors. To avoid contamination by XCR1^+^CADM1^+^ DC1, CADM1^+^ macrophages were sorted according to the gating strategy 2 which includes a lineage excluding lymphocytes and DCs (BTLA, CD26, CD3, CD19).

**Figure S3.**
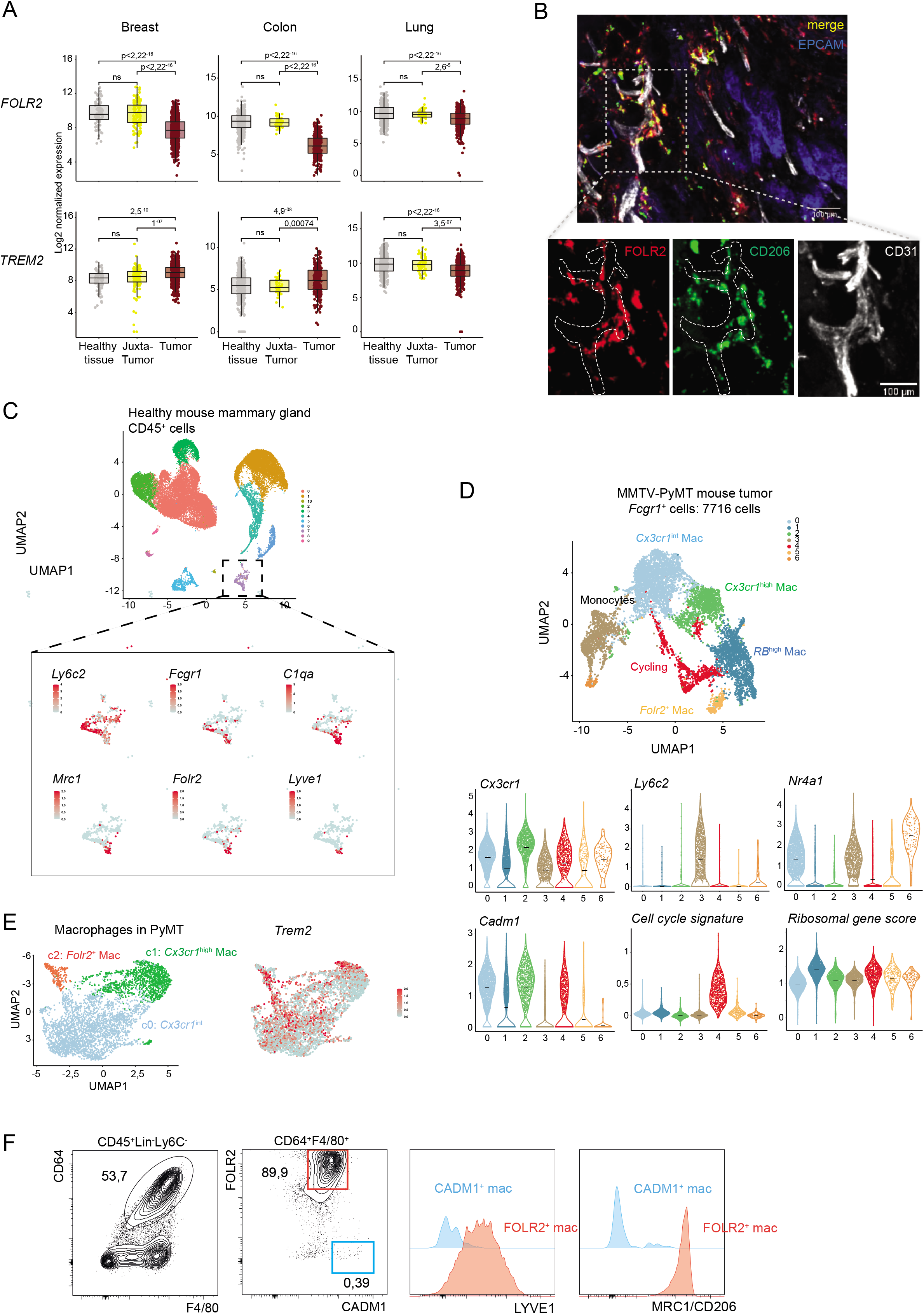
**A**. *FOLR2* and *TREM2* mRNA expression between non-disease healthy tissue, tumour-adjacent normal tissue, and tumour tissue from GTEx and TCGA datasets. **B**. Representative confocal immunofluorescence images performed in primary breast cancer tissues. showing FOLR2^+^CD206^+^ macrophages co-localizing with CD31^+^ vessels, near EPCAM^+^ tumor cells. **C**. UMAP plot vizualization of *Cd45^+^* hematopoietic cells isolated from healthy mouse mammary gland from the Han *et al*. dataset. Feature plots illustrating expression of *Ly6c2, Fcgr1, C1qa, Mrc1, Folr2, Lyve1* in macrophages. **D**. UMAP plot vizualization of *Fcgr1^+^* cells (n= 7716) isolated from mammary tumors of 23-week-old MMTV-PyMT mice (n=2). Violin plots illustrating expression distributions of *Cx3cr1, Ly6c2, Nr4a1, Cadm1*, a cell cycle gene signature and a ribosomal gene score. **E**. Feature plots illustrating expression of *Trem2* in macrophage subsets from mammary tumors (see Fig. 3E). **F**. Flow cytometry analysis of CD64^+^F4/80^+^ macrophage populations in healthy mouse mammary glands.

**Figure S4.**
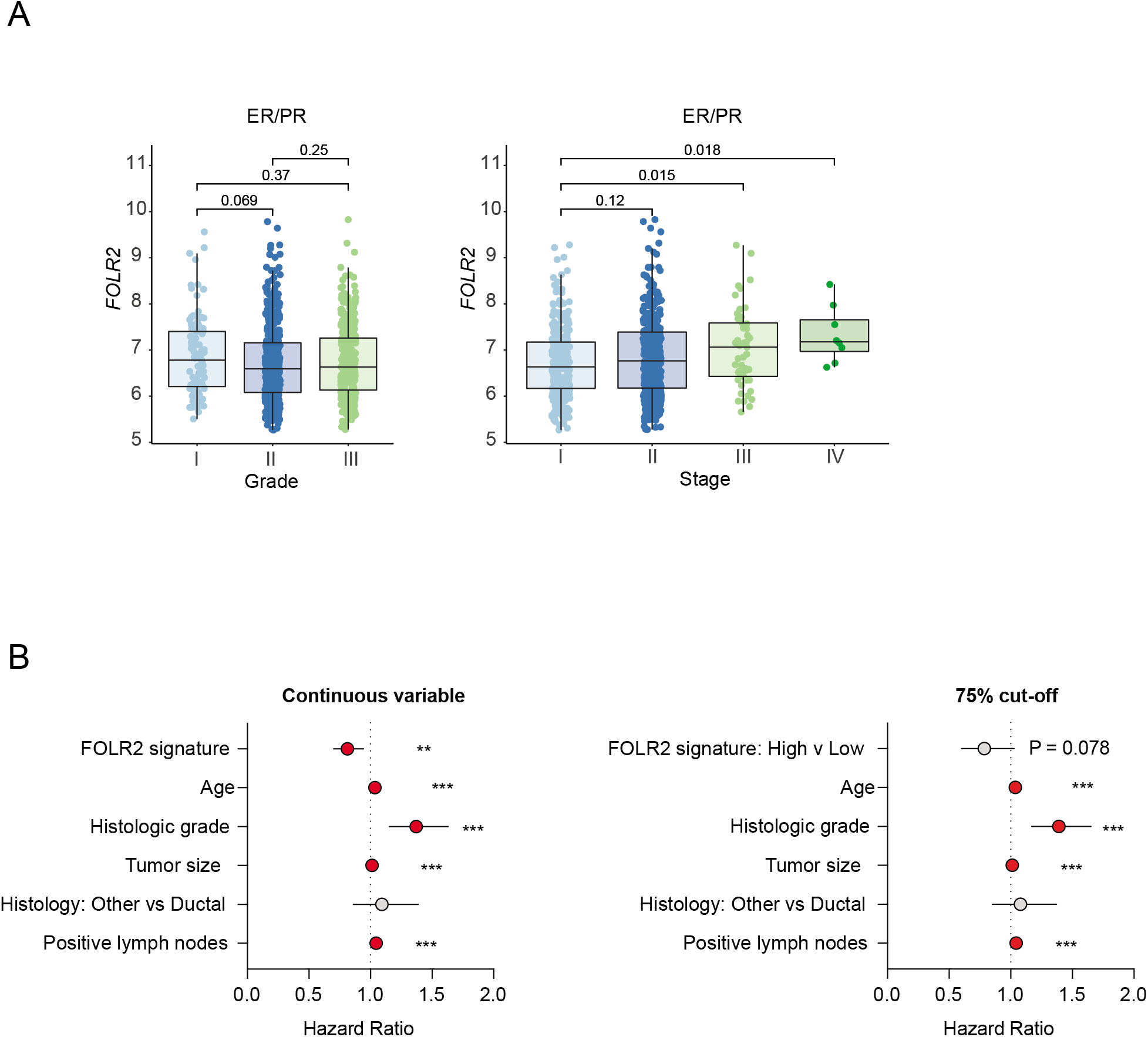
**A**. *FOLR2* mRNA expression in luminal (ER/PR) breast tumors of different grades and stages from the METABRIC dataset. **B**. Multivariate analysis of the prognostic value of the FOLR2 gene-signature as a continuous variable (left) or with a 75% cut-off, adjusted for age, histological grade, tumor size, histology (reference = ductal) and number of disease-positive lymph nodes. Hazard ratios with 95% confidence intervals are shown. Asterisks refer to P values from the Wald test for each variable.

**Figure S5.**
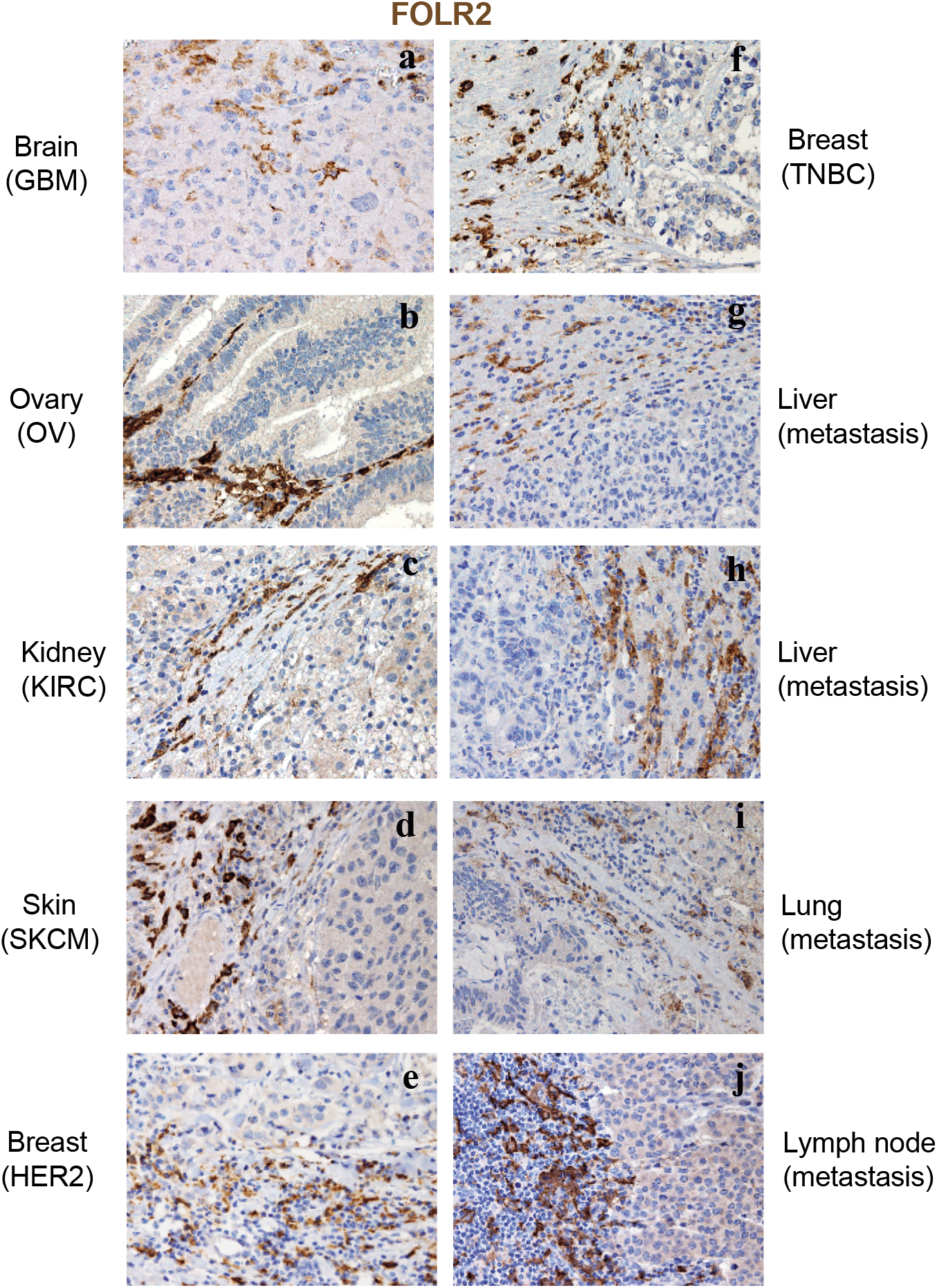
Spatial distribution of FOLR2^+^ TAMs in human primary and metastatic cancer. Sections are from cases of primary cancer including glioblastoma (**a**), serous ovarian carcinoma (**b**), renal clear cells carcinoma (**c**), cutaneous melanoma (**d**), HER2^+^ breast carcinoma (**e**), triple negative breast carcinoma (**f**), and cases of metastatic carcinomas to liver (**g-h**), lung (**i**) and lymph node (**j**), stained as labeled.

**Figure S6.**
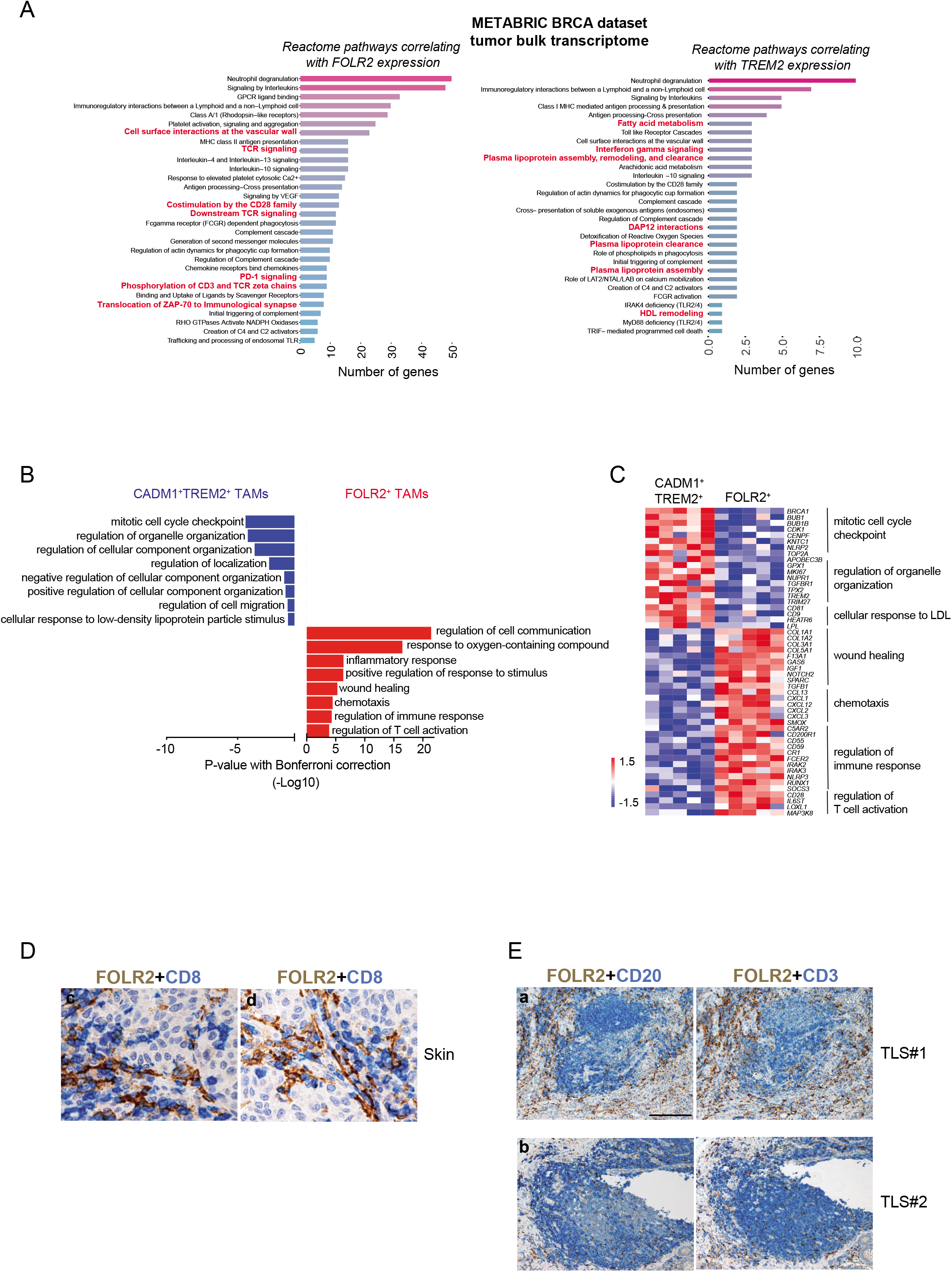
**A**. Bulk transcriptome of breast tumors from the METABRIC dataset. Reactome pathway analysis for genes positively correlating with either *FOLR2* or *TREM2* expression (r ≥ 0,4). P value < 0,01. **C-D.** Differential enrichment of gene pathways analyzed in bulk RNAseq of FOLR2^+^ and CADM1^+^TREM2^+^ macrophages isolated from primary tumors (see Fig 2H) (ClueGO and CluePedia Cytoscape apps). Heatmap showing expression by FOLR2^+^ and CADM1^+^TREM2^+^ macrophages of representative genes of each pathways. **E**. Sections are from cutaneous melanomas and stained for FOLR2 and CD8. Double stain illustrates the interaction of FOLR2^+^ TAM with CD8^+^ T-cell in melanomas. Magnification = 400X. Sections are counterstained with Mayer’s hematoxylin. **F**. Sections are from two colorectal carcinomas (**a, b**) and stained as labeled. FOLR2^+^ TAM are found at the periphery of tertiary lymphoid structure, defined by CD20 and CD3 aggregates, as well as within the CD3^+^ T-cell area. Magnification = 100X. Sections are counterstained with Mayer’s hematoxylin.

**Table S1**

Table of all patients involved in this study

**Table S2**

Differentially expressed genes for the clusters 0 to 5 from the scRNAseq of CD11c^+^HLA-DR^+^ cells isolated from primary breast tumors and metastatic LNs (relates to Fig. 1D).

**Table S3**

Z-scores of regulons inferred using the SCENIC pipeline for the APOE^+^ macrophages clusters found in scRNAseq (relates to Fig. 2C).

**Table S4**

Differentially expressed genes between macrophages clusters (relates to Fig. 2D).

**Table S5**

Z-scores for expression levels of selected human and mouse ortholog genes evolutionarily conserved across species (relates to Fig. 3F).

**Table S6**

Gene list relating to Fig. 4A

**Table S7**

List of tumor sections used for immunohistochemistry (relates to Fig. 3C, Fig. 5C-D, Fig. S5C, Fig. 6E-F, Fig. S6B-C).

**Table S8**

Gene signature used in the correlation dot plot Fig. 6A.

**Table S9**

List of antibodies used in the study

**Table S10**

Human and mouse gene symbols (relates to Fig. 3F-G).

